# Average Nucleotide Identity based *Staphylococcus aureus* strain grouping allows identification of strain-specific genes in the pangenome

**DOI:** 10.1101/2024.01.29.577756

**Authors:** Vishnu Raghuram, Robert A Petit, Zach Karol, Rohan Mehta, Daniel B. Weissman, Timothy D. Read

**Affiliations:** Microbiology and Molecular Genetics Program, Graduate Division of Biological and Biomedical Sciences, Laney Graduate School, Emory University, Atlanta, Georgia, USA; Division of Infectious Diseases, Department of Medicine, Emory University, Atlanta, Georgia, USA; Department of Physics, Emory University, Atlanta, Georgia, USA

## Abstract

*Staphylococcus aureus* causes both hospital and community acquired infections in humans worldwide. Due to the high incidence of infection *S. aureus* is also one of the most sampled and sequenced pathogens today, providing an outstanding resource to understand variation at the bacterial subspecies level. We processed and downsampled 83,383 public *S. aureus* Illumina whole genome shotgun sequences and 1,263 complete genomes to produce 7,954 representative substrains. Pairwise comparison of core gene Average Nucleotide Identity (ANI) revealed a natural boundary of 99.5% that could be used to define 145 distinct strains within the species. We found that intermediate frequency genes in the pangenome (present in 10-95% of genomes) could be divided into those closely linked to strain background (“strain-concentrated”) and those highly variable within strains (“strain-diffuse”). Non-core genes had different patterns of chromosome location; notably, strain-diffuse associated with prophages, strain-concentrated with the vSaβ genome island and rare genes (<10% frequency) concentrated near the origin of replication. Antibiotic genes were enriched in the strain-diffuse class, while virulence genes were distributed between strain-diffuse, strain-concentrated, core and rare classes. This study shows how different patterns of gene movement help create strains as distinct subspecies entities and provide insight into the diverse histories of important *S. aureus* functions.

**Importance:** We analyzed the genomic diversity of *Staphylococcus aureus*, a globally prevalent bacterial species that causes serious infections in humans. Our goal was to build a genetic picture of the different strains of *S. aureus* and which genes may be associated with them. We used a large public dataset (>84,000 genomes) that was re-processed and subsampled to remove redundancy. We found that individual genomes could be grouped into strains by sharing > 99.5% identical nucleotide sequence of the core part of their genome. We also showed that a portion of genes that are present in intermediate frequency in the species are strongly associated with some strains but completely absent from others, suggesting a role in strain-specificity. This work lays the foundation for understanding individual gene histories of the *S. aureus* species and also outlines strategies for processing large bacterial genomic datasets.

## Introduction

*S. aureus* is a ubiquitous human pathogen capable of causing numerous disease manifestations, including more than 100,000 bloodstream infections in 2017 in the US alone^1^. *S. aureus* genomes typically have a ~2.8 Mbase chromosome and zero to a few plasmids. Like other bacterial pathogens, its success at responding to pathogenic niches comes from both adaptations in the “core” portion of the genome and non-core genes that form the extended species genome, or “pangenome” ^2^. Non-core genes form part of the extensive genetic repertoire for evading the immune response and damaging the host and have allowed *S. aureus* to survive treatment with various antibiotics developed since the middle of the twentieth century ^3–6^.

Microbiologists have long known that there are consistent differences in phenotypes between taxonomic groups below the species level in *S. aureus*. Different “strains’’ have been shown to be more likely to cause specific disease etiologies than others. Examples are Multi-Locus Sequence Type (MLST) ST582, which is associated with scalded skin syndrome ^7^ and livestock associated CC97 infections ^8^. Among other phenotypes, strains also show different propensity to acquire drug resistance genes, high or low levels of toxin production, and can produce different spectra of mutations when under strong selection ^9–12^. Understanding the genetic basis of strain-specificity therefore offers potential insight into many mechanisms that define *S. aureus* pathology. Interest in strain-specificity has also been prompted by attempts to use shotgun metagenomic data to define environmental conditions that separate different genotypes with species ^13,14^. However, the cardinal problem with these approaches is that there is no generally accepted bacterial strain definition appropriate for the genomic era. Instead, the term “strain” has been used loosely to apply to different levels of sub-species variation.

The aims of this work were to seek a consistent definition of a *S. aureus* strain that could be applied to genomic and ultimately metagenomic data, to understand which portions of the non-core genome were strain-associated and to survey the extent of strain variation in the public data. We used an approach based on an earlier workflow ^12^, where we reprocessed all extant public Illumina whole genome shotgun (WGS) data. Here, we refined the strategy by implementing stringent steps to filter WGS potentially contaminated with other bacterial contigs and *S. aureus* mixtures. We also included high-quality complete genomes and dereplicated the final data set to remove very highly similar sequences. Critically, we opted to define relationships between genomes based on average nucleotide identity (ANI), rather than relying on the traditional clonal complex and sequence type designations of multi-locus sequence typing.

## Results

### ANI threshold of 99.5% defines 145 *S. aureus* strains from a large public genome dataset

To get a global view of *S. aureus* genetic diversity, we used all complete genomes without undefined (“N”) base calls and all Illumina whole genome data sets of the species available on the NCBI website in September 2022. The 83,383 whole genome data sets were filtered down to 58,034 (56,771 short read genomes + 1,263 complete genomes) based on having high sequence depth and quality, having no non-*S. aureus* genome content, and not being potential intraspecies mixtures based on minor-allele frequency (**Figure 1**, **Figure S1A; Methods**). To remove redundancy, the high-quality shotgun sets and 1,263 complete genomes were clustered based on a mash distance of 0.0005 (approximately 50 SNPs) ^12,15,16^. A randomly chosen representative of each of these 7,954 “substrains’’ was selected for downstream analysis.

**Figure 1:**
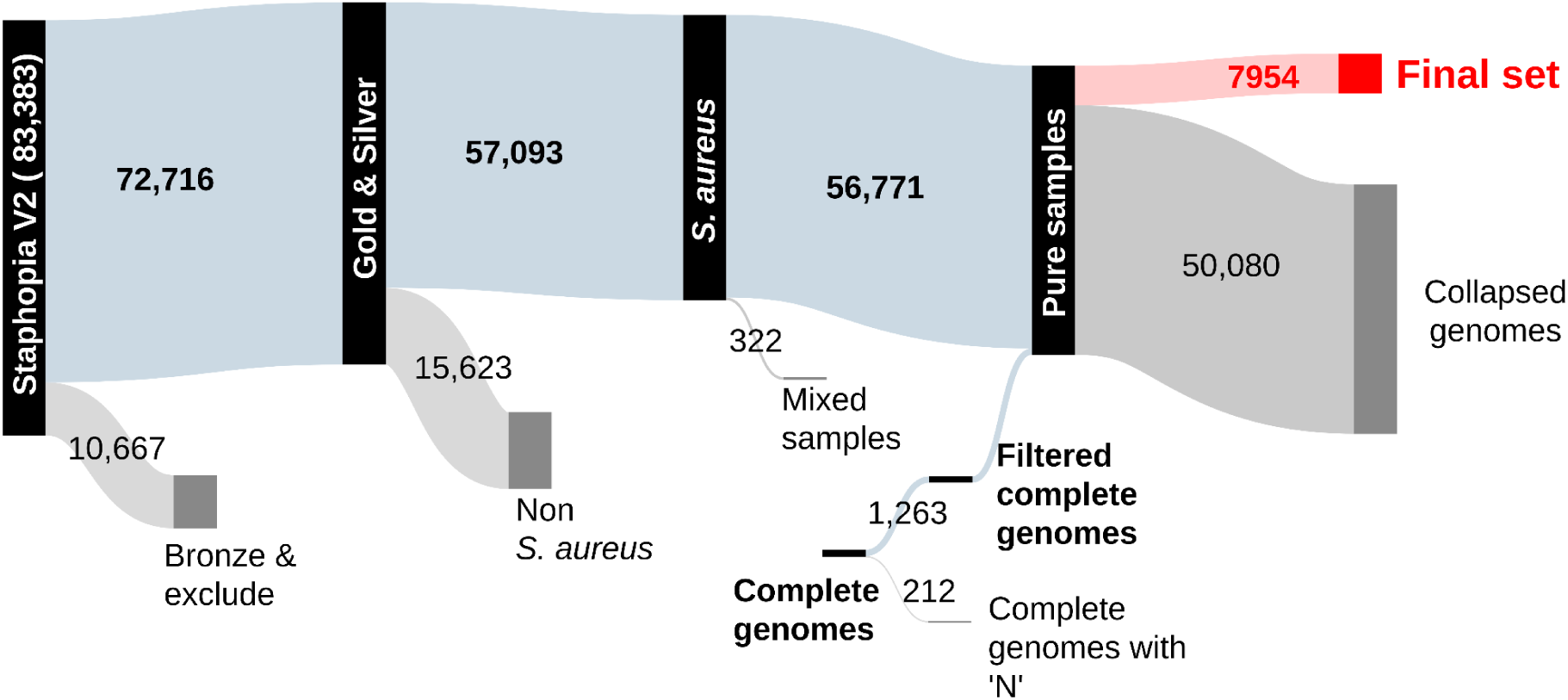
Sankey diagram showing the fate of 83,383 *S. aureus* whole genome shotgun datasets and 1475 complete genomes through processing and filtering.

The 7,954 representative substrains came from 1706 multi-locus sequence types (STs), with 386 substrains not belonging to a previously assigned ST. The uneven distribution of genomes across substrains and STs reflected the sampling skew towards well-known *S. aureus* strains from predominantly clinical settings. We found that the fifteen substrains that represented the most collapsed genomes, comprised 50% of the shotgun datasets. The most numerous substrain, from CC22, comprised 7688 of the 58,034 whole genomes (13%), while there were 5597 substrains represented by only one genome. 3857 out of 7,954 substrains(48%) were in ten most abundant STs (ST5, ST8, ST30, ST398, ST45, ST1, ST22, ST15, ST59 and ST239), representing 39,366 out of 56,771 genomes (69%).

The 7,954 representative substrains were used to create a species pangenome (the ‘7954-set’), using the PIRATE software^17^ based on a minimum 50% protein sequence identity. 9,533 distinct orthologous gene families were identified (we use the shortened “genes” to refer to these gene families in this manuscript). Of these genes 2,008 (21.1%) were considered core (found in > 95% of the genomes), 71.3% (6,794) were rare (<10% of genomes) and 7.7% (731) were intermediate between core and rare. 90% of genes were in single copy (**Figure S2**).

When pairwise average nucleotide identity (ANI) between substrains based on the concatenated nucleotide sequences of the core genes (2,101,692 nt) was plotted as a histogram there was a clear pattern of three strong peaks separated by distinct valleys (**Figure 2A**). The left peak (smallest AN distances), we interpreted as intra-strain distance, the second and third as between-strain distances within the two major *S. aureus* clades ^18^, and between the clades, respectively. The threshold for intra-strain relatedness appeared to be at, or very near to, 99.5%: identical to a value suggested by Rodriguez-R et al to separate strains across 330 bacterial species^19^. When we used 99.5% as a threshold for clustering we obtained 145 groups of genomes that we termed “strains’’ and marked each with a suffix “S99.5_”. All strain clusters had median within-cluster ANI > 99.7 (**Figure S1B**). Both gene discovery rate and lineage discovery rate were improved by dereplicating the initial 58,034 genomes compared to using a random set (**Figure S1C, S1D**).

**Figure 2:**
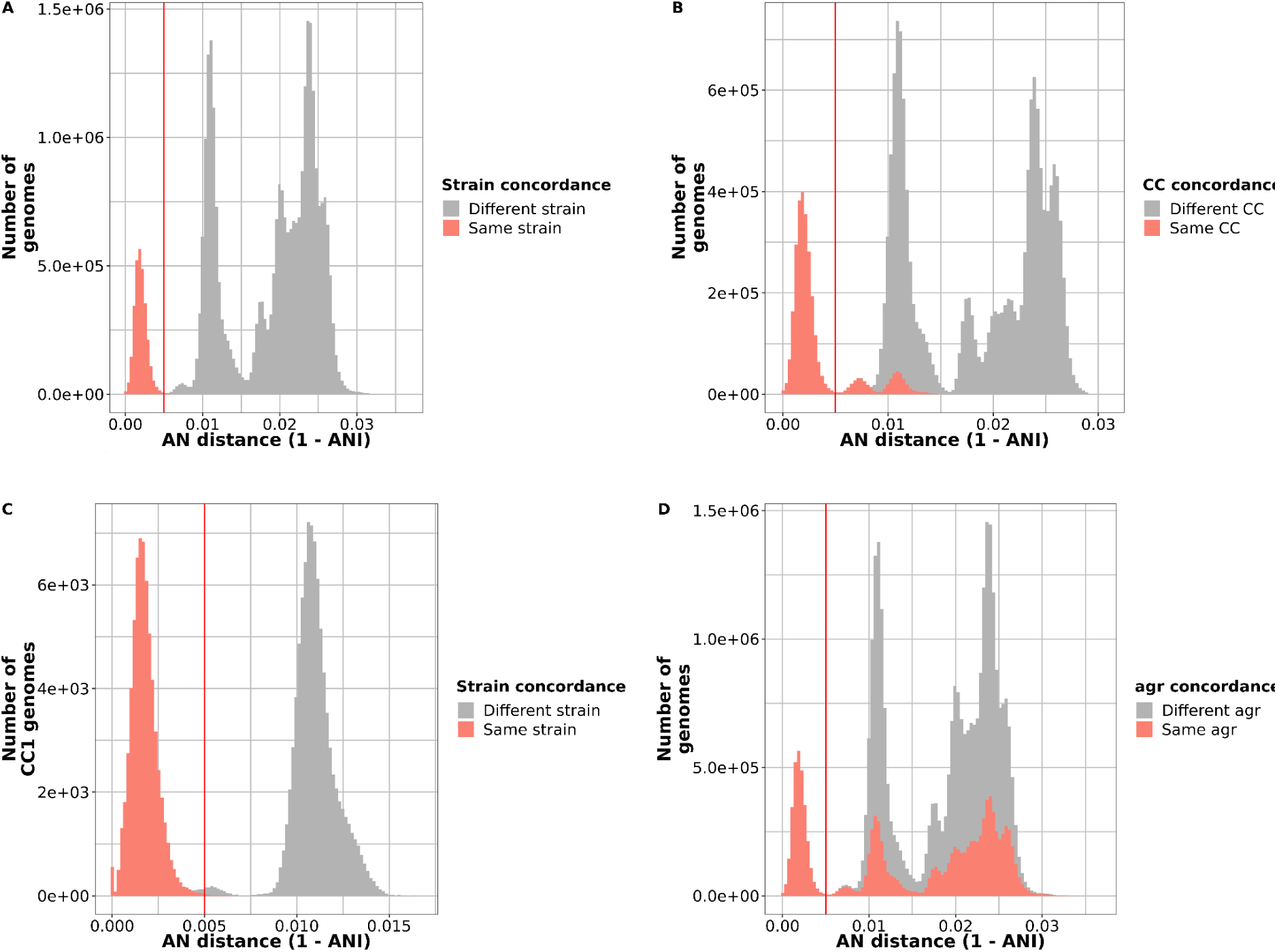
An average nucleotide identity of >99.5% in the core genome defines the strain boundary of *S. aureus*. For our dataset of 7954 substrains, all-vs-all pairwise Average Nucleotide (AN) distances were plotted as a histogram. (A) Sample pairs less than 0.005 AN distance apart (i.e. greater than 99.5% ANI) were grouped as a strain. (B) Strains and clonal complex designations don’t exactly overlap. The pairwise AN distance histogram was colored by whether the genomes were in the same clonal complex. (C) CC1 genomes are in different strains. AN distances of genomes assigned to CC1 showing that there are within- and between-strain distances. (D) Genomes in the same strain have the same agr group.The pairwise ANI distance histogram was colored by whether the genomes were in the same *agr* group.

Currently, ten clonal complexes (CCs) of closely related STs are defined by the *S. aureus* PubMLST site^20^. Of these, CC1, CC5, CC8, CC15, CC45, CC97 and CC121 were split into 14, 3, 6, 2, 3, 5 and 5 strains, respectively, at the 99.5% clustering threshold (**Figure 2B**). In the case of CC1, ten strains had 7 or fewer substrains (**Figure 2C**). Two strains, S99.5_9 and S99.5_36, contained substrains that had been assigned to different CCs. S99.5_36 had substrains assigned CC1, CC8 and CC97 (56, 3 and 1, respectively) and S99.5_9 had substrains from CC1 and CC97 (17 and 1, respectively). Substrains from different CCs assigned to both S99.5_9 and S99.5_36 had at least 5 alleles in common, suggesting that they were close to the threshold of being in the same CC by the rules of MLST assignment (which require 5/7 common alleles). Across all strains we found that >99.9% of genomes in the same strain had the same *agrD* specificity allele (1-4) of the *agr* quorum sensing system (**Figure 2D**). (The one exception was strain PS/BAC/317/16/W (GCF_018093225.1)^21^, the single *agr* group 2 genome in 4,469 CC30 genomes). This result confirmed an earlier genome-based screen^15^ showing that *agr* type is strongly strain specific in *S. aureus*.

We noted that there was a “bump” of pairwise distances (~99.5-99.1% ANI) in the otherwise clear gap between within-strain and between-strain comparisons (**Figure 2A**). When we clustered substrains at 99.1% core genome ANI we found that 30 99.5%-defined strains merged together to form 115 putative strains. One of the merged strains comprised genomes of S99.5_2 and S99.5_27, both largely mapped to CC8. The S99.5_27 strain consisted of ST239, which is known to have been created by recombination of a large portion of a CC30 genome with a CC8 background ^22,23^. The other 9 sets of merged strains consisted of a small number of genomes. For two of the merged strains, we had a complete genome which we used to align 10,000 bp sliding windows against a genome from the same strain at 99.5% ANI and one from a different strain that was merged at 99.9% ANI. These were strains S99.5_33 and S99.5_4 (both mapped to CC45) S99.5_7 and S99.5_111 (CC15), each pair merged into one strain using ANI 99.1% thresholds. Neither analysis revealed the clear pattern of large scale genome replacement seen in ST239.

### Intermediate frequency genes in the pangenome can be divided into strain-concentrated and strain-diffuse

We wanted to know what proportion of the *S. aureus* accessory gene was strongly linked to strain background, in the same manner as *agr* type. We adapted the commonly used genetic statistic F_ST_ (also known as fixation index) as a measure of segregation of a gene between different strains ^24^. F_ST_ of 0 indicated a gene that displays no genetic segregation, i.e it was indiscriminately found across different strains. In contrast, F_ST_ of 1 indicated perfect genetic segregation, with the gene limited to all members of a group of strains. Rare and core genes were constrained in their distribution and had uninformative F_ST_ scores around 0. Therefore we concentrated our analysis on intermediate gene families.

Strikingly, the F_ST_ distribution across intermediate genes showed a distinct bimodal distribution (**Figure 3A**). This pattern disappeared when the strain labels were randomly mixed and F_ST_ recalculated (**Figure 3B**), reverting to a normal distribution, showing that it was a feature of the specific population structure of *S. aureus* rather than an inherent property of the data. From this result we divided intermediated genes into two groups based on a F_ST_ threshold of 0.75. Those genes with high F_ST_ (296/731 (40%) intermediate genes), which we termed “strain-concentrated” were strongly linked to strain backgrounds, while those with low F_ST_ (“strain-diffuse”) (495/731 (60%) intermediate genes) were more promiscuous with respect the strain background. These patterns were illustrated using ten *S. aureus* toxins with a range of F_ST_ scores: Leukocidins LukFS (Panton Valentine Leukocidin) and LukED, Toxic Shock Syndrome toxin 1 (TSST), superantigen-like protein SSL8, and different types of *Staphylococcal* Enterotoxins (SEA, SEB, SEG, SEU) (**Figure 4**). Leukocidins comprise two proteins, the F component and the S component, both acting synergistically to form pores in host-cell membranes ^25^. TSST, SEs and SSL8 are superantigens or superantigen-like proteins (SAs), highly potent toxins that can elicit severe inflammatory responses and other immunomodulatory effects ^26^. The leukocidin LukFS, enterotoxins SEA & SEB, and TSST, showed high levels of gain and loss on the species tree typical of low-F_ST_. In contrast, the enterotoxins SEG and SEO, Leukocidin LukED, found together on genomic island vSaβ had high F_ST_ (> 0.9) and were either almost entirely present or absent in each strain background.

**Figure 3:**
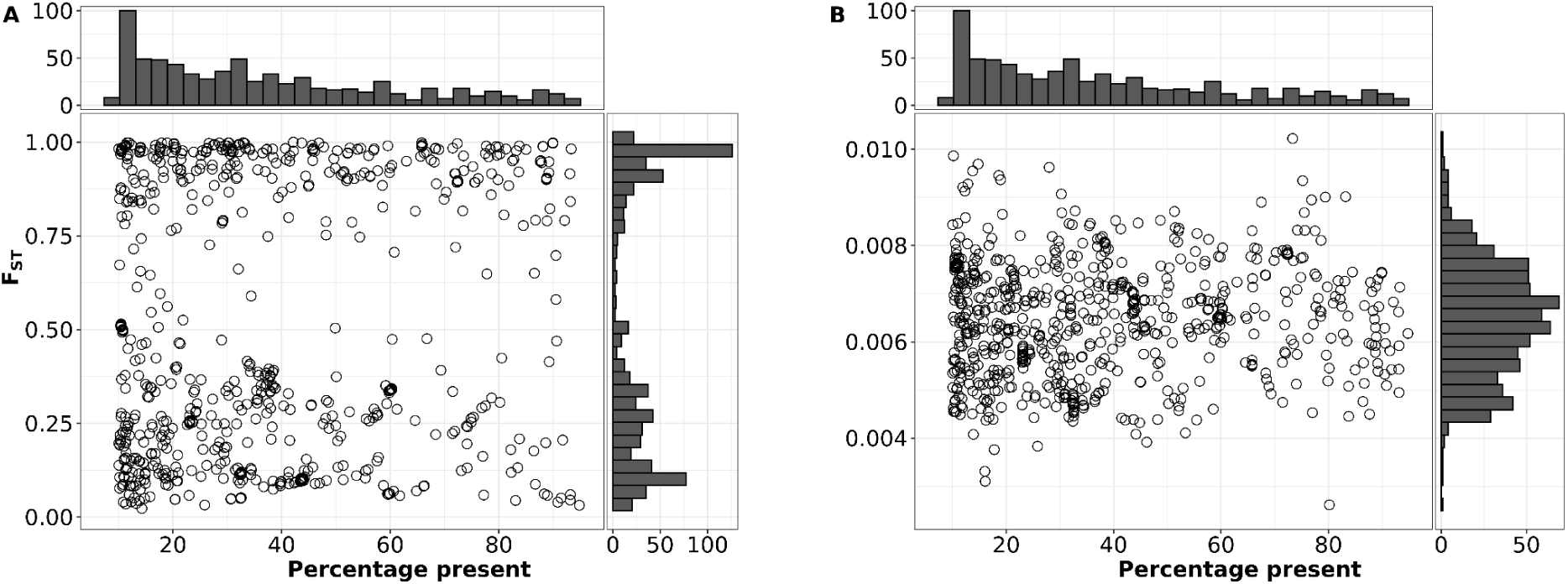
Bimodal distribution of F_ST_ for intermediate genes. Each circle represents an individual intermediate gene from the 7954 substrain pangenome. Percentage prevalence on the x axis is the percentage of genomes the gene is found in. F_ST_ or ‘fixation index’ is on the y axis. (A) F_ST_ scores calculated for each intermediate gene with 99.5% ANI-based clustering. (B) As a control, F_ST_ scores were calculated for each intermediate gene when clusters were randomly assigned.

**Figure 4:**
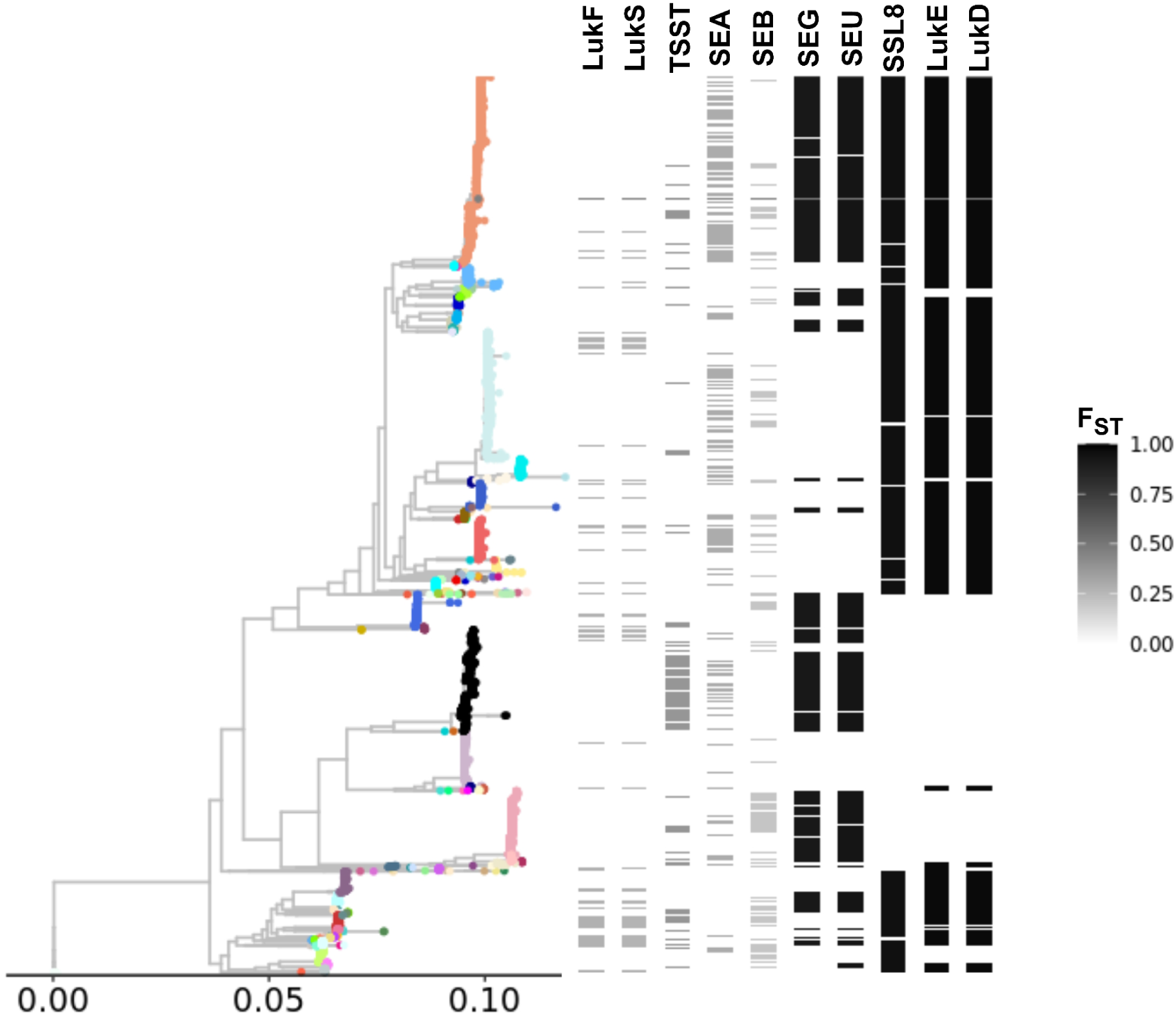
Strain-group specificity and co-occurrence of specific Staphylococcal toxins. Core genome of the 7954-set. Heatmap on right shows presence absence and FST of specific Staphylococcal toxins - Panton-Valentine Leukocidin (LukF and LukS), Toxic Shock Syndrome Toxin (TSST), and Staphylococcal Enterotoxins type A, B, G, U (SEA, SEB, SEG, SEU), Superantigen like protein (SSL8), Leukocidin ED (LukE, LukD). The colors of the whole-genome phylogeny are based on strain assignments.

We also used F_ST_ to test whether there was any association between the *agr* type of a strain and intermediate gene distribution but found no similar pattern (**Figure S3**).

To investigate the differences between strain-concentrated and strain-diffuse genes further in a *S. aureus* pangenome with more balanced sampling, we created the “740-set”, created by randomly sampling 20 shotgun assembled substrains from the most common 37 strains. The 740-set had similar numbers of core and intermediate genes (2,139 and 739, respectively) to the 7954-set but fewer rare genes (2,687), the latter expected to increase with the number of genomes sampled in a species. The F_ST_ distribution of the 740-set to the original pangenome was almost identical.

When we plotted the number of strains each gene was found in given the numbers of genomes we saw two distinct patterns. The strain-concentrated genes were close to the minimum possible number of strains for a given gene (dashed red line), while the strain-diffuse genes were more similar to the shape of a random assortment of strains (asymptotic exponential distribution; dashed blue line)(**Figure 5A**). Strain-diffuse genes were present in markedly more strains at a given prevalence than strain-concentrated From **Figure 5A** it was clear that rare gene distributions were extensions of the trends seen in intermediate genes. These trends could not be discerned in the 7954-set because the number of substrains represented in each strain was unbalanced.

**Figure 5:**
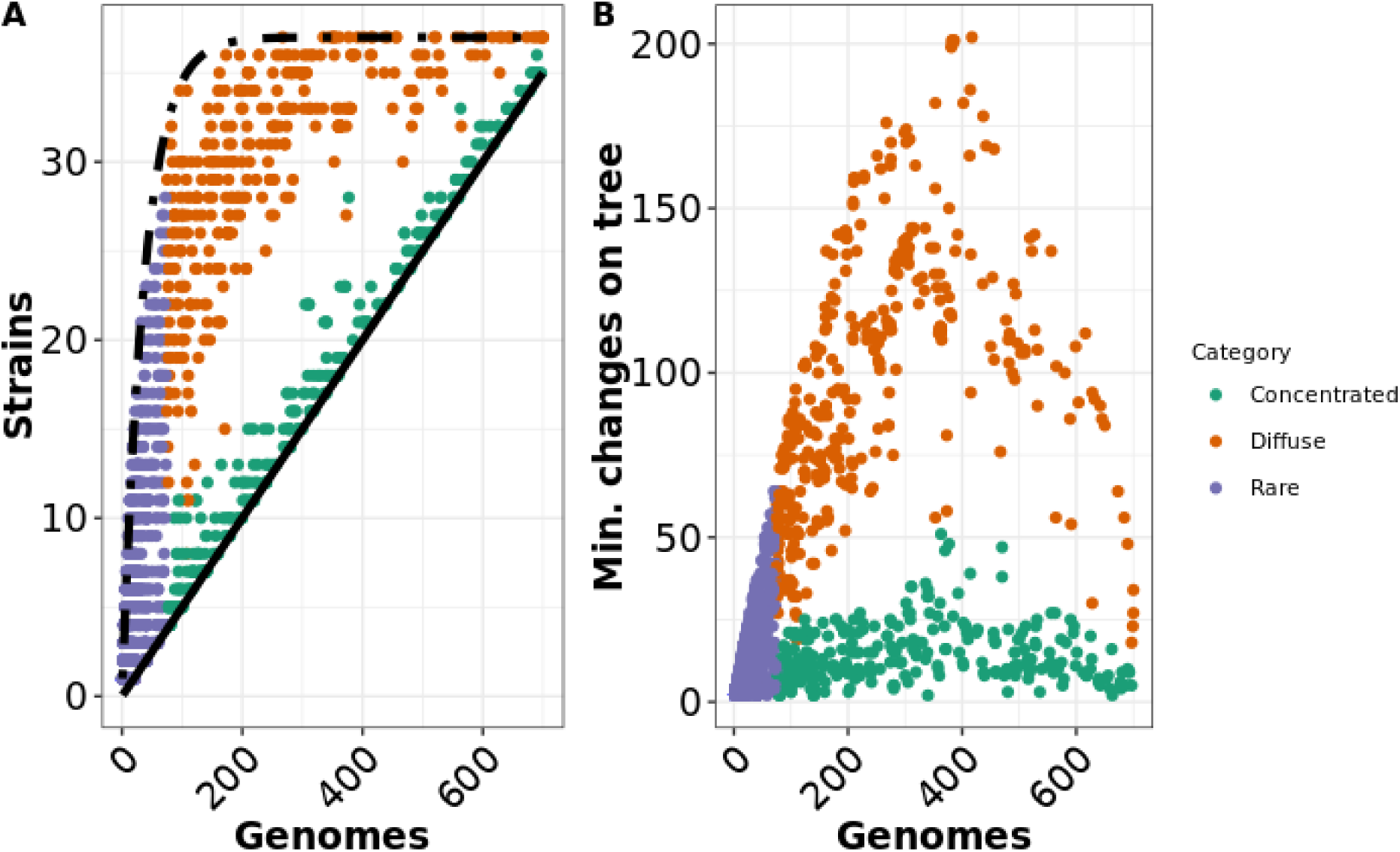
Relationship between gene prevalence, number of strains and homoplasy for non-core genes. Each dot represents a non-core gene in the 740-set pangenome. Purple = rare genes, green = concentrated, Brown = diffuse. In panel a, the relationship between overall prevalence (number of genomes out of 740) and number of strains (out of 37) each gene is found in is shown. The curves for the theoretical minimum number of strains for a given number of genomes (x/20) is shown in solid black and the extreme random distribution (37*(1-exp(-x/37)) is shown in dashed black. Panel b shows the relationship between prevalence of estimated number of changes on the species tree calculated by homoplasyfinder^27^.

**Figure 3 and 4** depict a pattern where strain-diffuse genes appeared to undergo gain and loss on the phylogenetic tree at a higher rate than strain-concentrated genes. Based on the results of homoplasyFinder^27^ analysis on genes arrayed on the core gene phylogeny of the 740-set, we found this pattern was consistent across all intermediate genes (**Figure 5B**). Strain-concentrated genes mostly had fewer than 30 minimum predicted state changes on the tree and there was no trend in increase of this number with prevalence. Strain-diffuse genes had a higher rate of character state change, which rose with prevalence initially but fell with the most common genes, probably due to saturation of available state changes.

Because of the relatively slower rate of gene gains and losses, the strain-concentrated genes contributed more to characteristic strain-specific differences in gene content than strain-diffuse genes. This could be effectively visualized using t-SNE (t-distributed stochastic neighbor embedding; **Figure 6**). When strain-concentrated was used as input for t-SNE, the genomes that comprised individual strains were resolved into distinct spatial units (**Figure 6C**). However, there was no similar pattern when strain-diffuse was used (**Figure 6B**). Rare genes produced an intermediate result, with some distinctive strains and some areas of the plot with mixtures of strains (**Figure 6A**). When all non-core genes were used the strains could be readily distinguished, indicating that for the t-SNE approach, the strain-specific structure of strain-concentrated and rare gene content was dominant to the non-strain specific strain-diffuse genes (**Figure 6D**). We also visualized the effect of the different classes of non-core gene is a way that was independent of strain classification: plotting the gene content similarity (represented by hamming distance) of each pair of genomes against the patristic distance on the core gene phylogeny (**Figure S4**). The rare and strain-diffuse genes had greater numbers of gene differences between strains very closely related to each other (Patristic distance < 0.005) but the rate of growth of the distance in strain-concentrated genes over larger distances on the phylogeny was greater. Together these results showed that strain-concentrated genes provided more information about gene content differences between strains than other non-core genes. We suspected that the underlying differences between the two groups of genes were due to strain-concentrated genes being primarily located on the chromosome and primarily spread between strains by homologous recombination, whereas strain-diffuse genes were on mobile elements such as prophages, plasmids and integrative conjugative elements that would be located more frequently on non-chromosomal contigs. This was supported by the rate of linkage to single copy highly conserved core genes (defined as whether the gene was found to be on the same contig) was much lower in strain-diffuse genes (65.5%) than strain-concentrated (86.5%). By comparison, the rates for rare genes were 61.5% and randomly selected genes were 93.5%. We used the geNomad software and database of mobile element gene ^28^ to see if there were different distributions in the different classes of genes in the pangenome. While differences between the classes were mostly statistically significant at p < 0.05 in pairwise Tukey’s tests (**Figure S5**), the difference in mean scores were mostly quite small, probably reflecting the relatively small size of the *S. aureus* training set for the software compared to our large pangenome sampling. The strain-diffuse genes had the most distinctive signal, having the lowest mean scores for “chromosome” and “plasmid” and highest for “virus”. This result corroborated the association of strain-diffuse genes with prophage regions of the genome.

**Figure 6:**
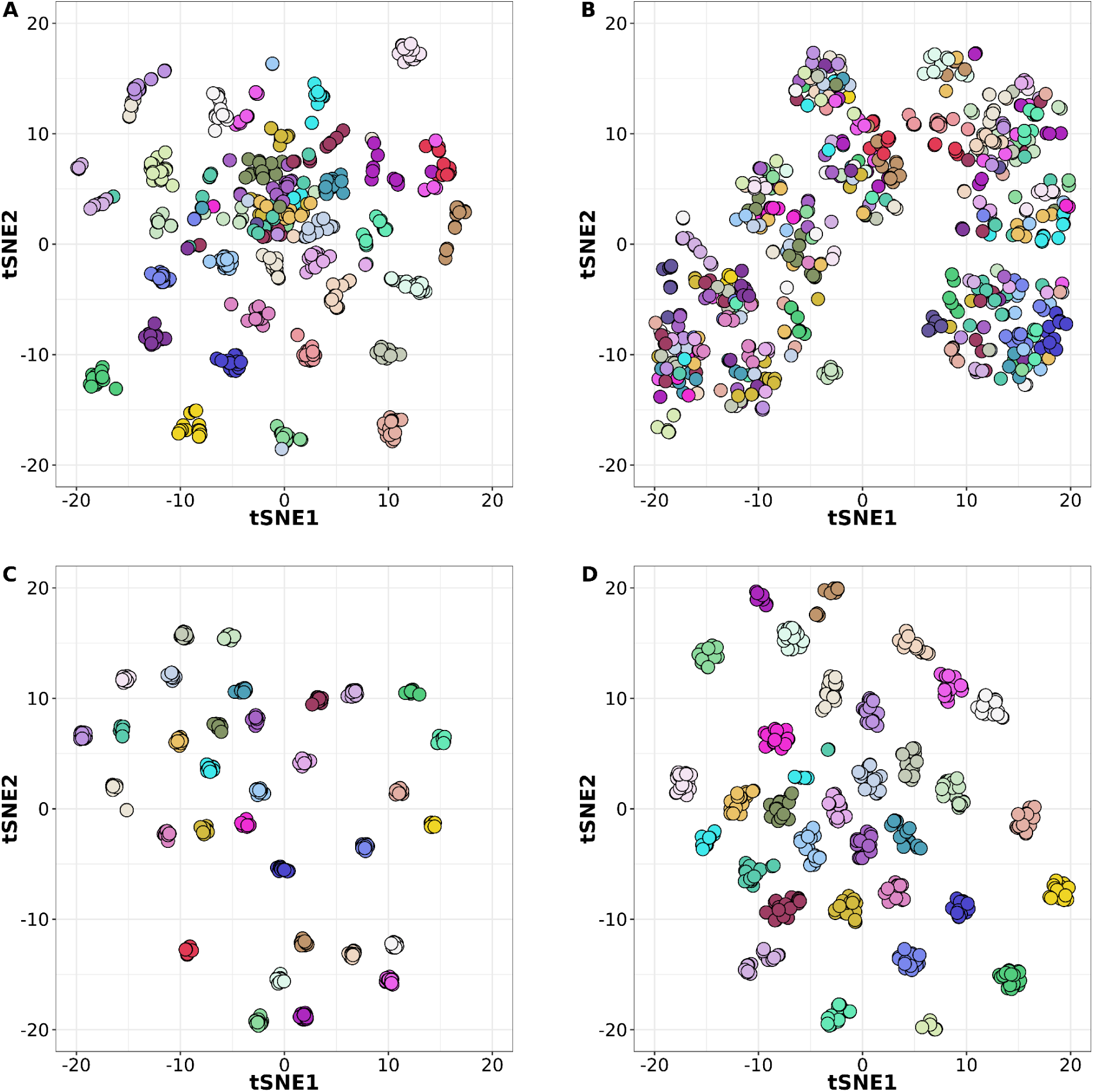
t-SNE analysis of 740-seq differentiated by non-core gene sets. Each dot represents one of the genomes of the 740-set colored by its strain membership. Different sets of non-core genes were used as input for the t-SNE: a) only rare; b) only strain-diffuse; c) only strain-concentrated; d) all non-core.

We noted that the intermediate genes had a lower median clustering threshold than the rare or core genes (the PIRATE software uses iterative thresholds at increasing stringency to find the final clustering threshold for a gene ^17^). To ensure the patterns seen were not an artifact of lower clustering, we ran the 740-set pangenome with a minimum clustering threshold of 90% amino acid identity (which we called “740-set-90”). While the more stringent clustering split several rare and intermediate gene families (the “740-set-90” pangenome consisted of 4,490 rare, 982 intermediate and 2,085 core) the characteristic divergence in features between strain-concentrated and strain-diffuse genes did not change (**Figure S6**). We also obtained similar results when the same analyses were run with the original 7,954 substrain pangenome, although the unbalanced nature of the collection (some strains had thousands of genomes, many only one) obscured the differences between strain-concentrated and strain-diffuse in regards the relationship between strains each gene was detected in at different prevalence (**Figure S6A**). The strain-concentrated genes though had many fewer predicted state changes on the phylogenetic tree (**Figure S6B**).

### Different non-core gene classes cluster in specific regions of the *S. aureus* chromosome, with a strong tendency for rare genes to be near the origin of replication

We used two orthologous methods to view the distribution of non-core genes on the *S. aureus* chromosome (**Figure 7**, **Figure S7**). In the first method we plotted the start coordinate of genes from 337 complete chromosomes(**Figure 7A**, **Figure S7**). There was noise in the exact coordinates of individual genes but overall this method showed discrete peaks in the locations of rare, and strain-concentrated and diffuse genes. The second method was to link non-core genes from all 7,954 substrains to the nearest core gene on the same contig (non-core genes on contigs without core genes were excluded). The gross patterns of distribution of the counts of non-core genes mapped to the core nearest core gene coordinate (**Figure 7B**) were similar to that in **Figure 7A**. Differences between plots in the proportion of genes within each category at each genomic bin (y-axis) were probably due to a combination of the indirect measurement of gene position in the linked core gene method and the fact that the 7,954 substrains were are more balanced reflection of *S. aureus* diversity than the 337 complete genomes.

**Figure 7:**
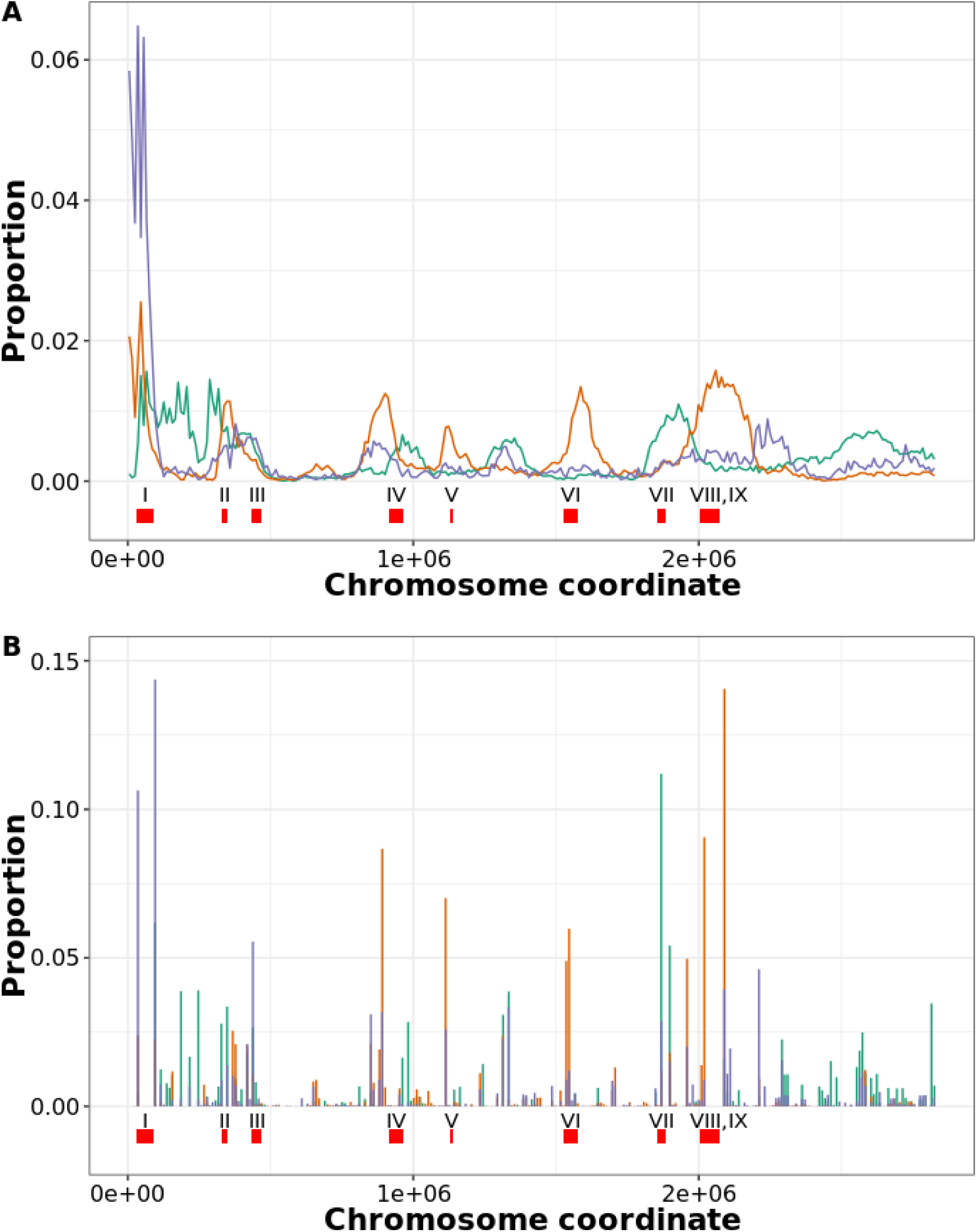
Distribution of different categories on non-core genes on the *S. aureus* chromosome using two orthologous methods. A: Location based on 337 complete genome sequences. The start site for every gene in each category was obtained for 337 chromosomes. The totals were placed in 10,000 bp bins on the chromosome and the proportion of the total for each class is plotted (i.e. the sum of the values of the 10,000 bins is 1). Purple = rare genes; green = strain-concentrated; brown = strain-diffuse. B: Location based on the nearest core gene. For all 7,954 substrains, the closest core gene on the same contig was determined. The x axis are start sites for the core genes of genome N315 (GCA_000009645)^29^. The values were binned and proportionalized as in A. For both A and B the location of selected features is shown: I = SCCmec; II = type VII secretion system; III = vSaα; IV = phiSa1; V=vSaγ; VI = phiSa2; VII = vSaβ; VIII = phiSa3; IX=vSa4). N315 coordinates are based on Gill et al ^29^ and Warne et al ^30^, except phiSa2 and phiSa3, which are from Mu50 and MW2, respectively.

Strain-diffuse and strain-concentrated genes had markedly distinct distributions on the chromosome and were mostly located as part of distinct clusters (**Figure 7**). This could also be seen clearly in the individual chromosomes of six substrains chosen to represent both MRSA and MSSA from three strains (**Figure S7**). The vSaβ genome island was a notably strain-concentrated-rich gene cluster, while the vSaγ island, phiSa2 and phiSa3 prophage were rich in strain-diffuse. The presence of strain-diffuse gene clusters was more variable between genomes than strain-concentrated clusters (**Figure S7**). Some genetic elements (e.g SCCmec, type VII secretion loci, phiSa1) contained a relatively high proportion of both types of intermediate genes. Three regions of the chromosome relatively rich in strain-concentrated genes (at approximate coordinates 100,00-300,000, 1,250,000-1,500,000 and 2,500,000-2,800,000) did not correspond to known genetic elements, although the first region contained several genes involved in polysaccharide capsule synthesis.

The high number of rare gene genes in the 0-100,000 region (which includes the SCCmec cassette) was an outlier compared to other chromosomal regions (p-value < 2.2e-16, Grubbs 1-tailed test) (**Figure 7**, **Figure S7**). This was the case in both MRSA and MSSA strains, suggesting that this region might be a hotspot for insertion of rare genes, possibly through plasmid integration, rather than being specifically linked to SCCmec.

### Functional differences in strain-concentrated and strain-diffuse genes

F_ST_ and prevalence of intermediate gene families can provide insight into ongoing evolutionary processes in the species. This is illustrated by analysis of three classes of genes encoding AMR (antimicrobial resistance), phage defense and virulence determinants (**Figure 8**). No AMR genes^31^ were found to be in the strain-concentrated group but were either rare or strain-diffuse (70 (82.4%) and 15 (17.6%), respectively). This result follows from the recent introduction of many AMR genes into *S. aureus* on mobile genetic elements and their frequent gains and losses below the strain level ^32^. The absence of fixation within strains also suggested possible loss of mobile elements in the absence of antibiotic selection. Genes associated with protection from phage infection in the defense-finder database ^33^ were mostly low prevalence (69/80 (86.3%) were rare and 10/80 (9.1%) intermediate had prevalence < 0.5). The low prevalence may reflect diversifying selection caused by phage countermeasures. However, unlike AMR genes, the majority of intermediate genes in this class were strain-concentrated, suggesting that defense from phage infection may help define *S. aureus* strains. Intermediate virulence genes (mostly toxins ^34,35^) in the AMRFinder+ database fell into two groups: one strain-diffuse with low prevalence and the other strain-concentrated with mostly higher prevalence. strain-diffuse virulence genes were mostly associated with prophages and Sa-PIs, while strain-concentrated genes were associated with the vSaβ genome island. This partition suggested an as-yet unexplained complexity in the hierarchy of functions that make up the toxin profile of an individual substrain.

**Figure 8:**
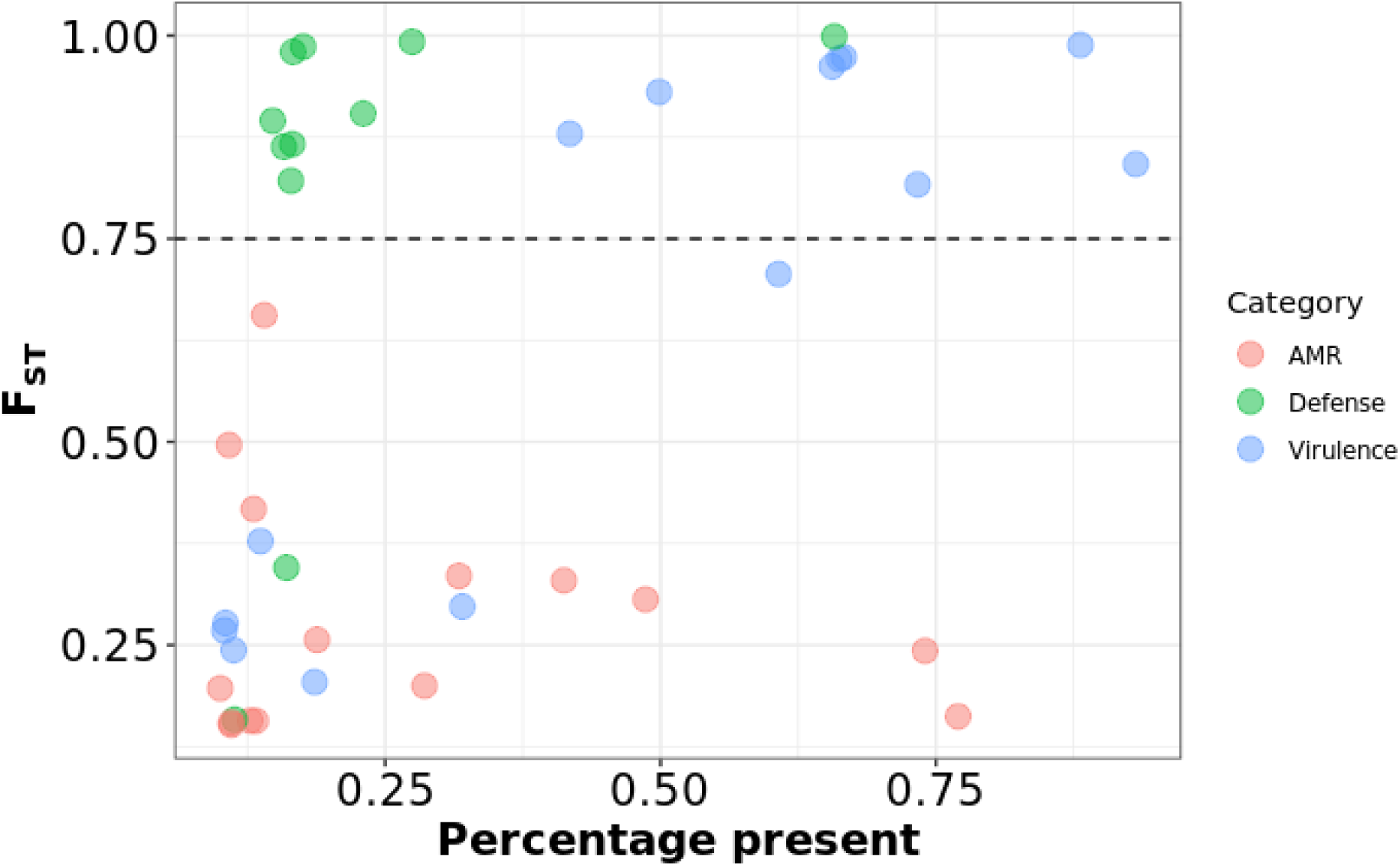
Prevalence versus Fst for intermediate antimicrobial-resistance (AMR), virulence and phage defense genes. AMR and virulence genes were identified using AMRFinder+^31^, phage defense genes were identified using defense-finder^33^. The dashed horizontal line represents the boundary between strain-diffuse and strain-concentrated.

## Discussion

In this study, we distilled a starting set of >84,000 *S. aureus* genome sequences to 145 strains using an ANI cutoff of 99.5%, which we found to be in a natural valley between clustered isolates. This threshold, or values close to it, has been reported in other studies as a bacterial subspecies boundary ^19^. A large number of *S. aureus* strains were rare (92/145 (63.4%) represented by 1-2 substrains). While this could represent some aspect of the true distribution of strain abundances in the species, it could also be a function of uneven sampling of *S. aureus* genomes. There are large ascertainment biases in selection as most strains are from clinical settings in western countries. It is probable that the number of strains will grow significantly in the future as we extend sampling.

There is no agreed term for the highest-level bacterial subspecies level although some names such as “genomevar” have been proposed ^19^. We had two reasons for choosing to use “strain”, which is a word frequently used in microbiology but currently has a multitude of different meanings. The first is to use “strain” in a way that gives it a precise definition, in this case genomes that cluster together above the natural 99.5% ANI gap. The second reason is that as the word is now frequently being used in metagenomic studies ^13,14,36,37^, and by choosing “strain” to mean the highest level of subspecies, this reduces the number of reference genomes needed to represent strain diversity in a species. This also increases the chances of discrimination between strains using the low coverage sequence read data often found in metagenome projects. However, sub-species terminology needs to be formalized through standards developed by consultation with the international microbiology community.

The 145 representative genomes defined here could be used for assignment of a new genome to an existing strain using fastANI or similar software. If the genome was found not to have >99.5% ANI to an existing strain it would be a candidate for a new strain. This simple approach for strain assignment has the advantage of not needing a core phylogeny calculated that is inherent to tree-based clustering and may turn out to be similarly accurate owing to the population structure of the within- and between-strain differences in the species (**Figure 1**). The existing MLST clonal complexes were mostly mapped with a 1:1 relationship to the strains defined, and the names, which are familiar in the literature, could be used as aliases for the strains. However, in some cases different genome backgrounds had been designated as part of the same CC but were split into more natural strain clusters by ANI. This is not surprising, as MLST schema was developed for PCR amplification and sequencing, before routine whole genome sequencing was available, and the seven loci used for assignment only cover a small portion of the variation in the chromosome ^38,39^. MLST, though useful for rapid strain typing, is outperformed by whole-genome based methods for lineage assignment ^39,40^.

Several pangenome studies with *S. aureus* genomes have been performed for epidemiological investigations ^41–46^, vaccine candidate discovery ^47,48^, and evolutionary phylogenomics ^49–52^. These produced a wide range of results, from 4,250 - 21,358 gene total pangenome size, with cores ranging from 890 to 2,700 genes(**Table S1**). The variability is a feature of the many factors that influence pangenome estimation, which can be classed into three main groups: sample collection, data quality and bioinformatics approaches. In terms of the collection, more individual genomes of a species tend to produce a larger number of gene families (in an “open” pangenome) and smaller core ^53^. Similarly, the more genetic diversity within the species increases pangenome size. We used essentially all the genome data available in the public domain by Fall 2022 (although we ended up excluding several thousand experiments based on quality (**Figure 1**). Therefore this study probably has the largest and most diverse input *S. aureus* set used to date. By reducing genome redundancy we also mitigated some of the overcounting of highly sampled clones in the public databases. Ideally, all genomes for a pangenome should be high-quality and complete. However, we chose to include shotgun assembled genomes, which may contain a certain percentage of missing genes due to contig breaks, to maximize diversity. Using shotgun assemblies also allowed us to sample multiple genomes from a larger number of strains, which was important for characterizing strain-diffuse and strain-concentrated genes. By reprocessing the data from raw reads, we were able to filter out lower quality data and have consistent assemblies (**Figure 1**). In tests, we found that pangenomes based on our shotgun assemblies produce similar metrics to those estimated used only complete genomes, as evidenced by the 740-set, which was composed entirely of shotgun data. For most complete genomes there is no matching raw read data available in public archives, so it is not possible to know whether the sequence is based on highly redundant reads coverage, as it is for our Bactopia processed genomes used here. The final group of factors concerns choices about bioinformatic software, and what parameters to use. Out of a wide range of open source options available we chose to use highly-cited tools Bakta ^54^ (which uses the Prodigal ^55^ gene finder) for annotation and PIRATE ^17^ for pangenome estimation. PIRATE iteratively increases the threshold to report the maximum identity that clusters each gene family and therefore avoids over-splitting gene families. PIRATE also identifies alleles within families without creating artificial paralog gene families. Tools that split paralogs into separate gene families (e.g ROARY ^56^ using default parameters) will also produce larger numbers of gene families and fewer core genes. The choice of minimum threshold for clustering proteins or genes into orthologous families (usually based on percentage identity of a pairwise alignment) is important. We realized from constructing the pangenome with a minimum 50% threshold that 85% of *S. aureus* genes families were clustered with at least the 90% identity. When we tested the 740-set pangenome with the minimum threshold increased to 90% we found a similar number of core genes (2139 at 50% minimum versus 2085 at 90% minium) but the number of non-core genes increased to from 3,426 to 5,472 (90%). This was because many intermediate gene families had been split at the higher threshold. However, the different threshold did not affect the key result of this study was that intermediate genes could be placed into two groups based on segregation with the strains defined by ANI using the F_ST_ statistic. Although we did not thoroughly explore different options in this study, pangenome estimation in *S. aureus* could be further optimized in future benchmarking studies based on the genome data collected here.

We defined three classes of *S. aureus* non-core genes with different properties. Strain-diffuse genes are maintained in the population yet have a high turnover, i.e they are gained and lost frequently (e.g LukFS, TSST, SEA, SEB in **Figure 4**). These genes are associated with mobile elements on the chromosome such as prophages, SaPIs and SCCmec and also often found on contigs unlinked to core genes, as would be expected of plasmids. These genes include niche-specific functions under high selection such as antibiotic resistance and certain toxins, which are classically segregated onto genetic elements that undergo frequent horizontal gene transfer in bacteria. *S. aureus* strain-diffuse genes are strikingly promiscuous in their strain background. Outside intra-strain comparisons, there is almost no signal of phylogenetic relatedness in strain-diffuse gene composition (**Figure S4**). This suggests high rates of horizontal transfer and, over the longer term, relatively weak barriers to genetic exchange compared to the strength of selection for strain-diffuse genes.

The second, previously unrecognized group of intermediate genes in *S. aureus* had a high F_ST_ score, indicating that they segregated closely with strain core gene background. Many of the genes cluster in the *S. aureus* genome islands, particularly vSaβ. The elements have been described as having complex, strain-specific genetic structure ^57,58^. Strain-concentrated genes also include significant virulence related functions located outside of previously defined genetic elements such as certain type VII secretion and capsule genes. strain-concentrated genes have many fewer predicted gene gains and losses than strain-diffuse genes (**Figure 5**) and a much stronger phylogenetic signal (**Figure S4**). This suggests that the rate of horizontal transfer of strain-diffuse genes is much higher and the probable reason is that they are on self-transmissible elements such as phages, plasmids (conjugative and mobilizable). The genome islands appear to have evolved from prophage or SaPIs that have acquired null mutations in their genes for site-specific recombination. We propose the mechanism of horizontal transfer of strain-diffuse genes is indirect: homologous recombination following introduction of DNA into the donor cell. Transduction is the dominant mechanism of DNA transfer in *S. aureus* and hence the genes likely rely on phages and/or SaPIs for their mobility.

Rare genes probably have properties either of strain-diffuse genes (high rates of HGT) or strain-concentrated genes (lower HGT rate) (**Figure 5**) but their low abundance makes calculation of F_ST_ the statistic meaningless. In other species (e.g *E. coli* ^59^) rare genes (and in some cases intermediate genes) have been reported to be strain-specific. We found that rare genes had strain-specificity levels between the two classes of intermediate frequency genes. In **Figure 3** some of the rare genes present in less than 10% of genomes are found in a significant majority 29/37 (78%) of strains. Both rare and strain-diffuse genes were frequently found to be genetically linked to core genes on the chromosome. While a higher proportion of strain-diffuse genes were distributed to a limited number of loci, representing common insertion points for SaPIs and prophages, it was a compelling finding of this study that a much higher proportion of rare genes were inserted in the region near the origins of transfer (approximate coordinates 1-100,000 in **Figure 7**). This was true in both MRSA and MSSA strains, hence the SCCmec element, which also integrates in this region, was not solely responsible for this pattern. This region of the chromosome, which is less dense in core genes, may serve as a “plasticity zone” ^60,61^ in *S. aureus* for capture of novel genes entering the species by HGT.

This study raises two questions about the manner in which the *S. aureus* genome evolves and the underlying selective pressures that drive the observed patterns: 1) what are the forces that create the “valley” of ANI in the range of 99.1-99.5% (**Figure 1**)? And 2) what are the functional implications of the partitioning of intermediate genes in strain-concentrated and strain-diffuse groups? The ANI valley implies that there is a limited time that strains can survive as coherent taxonomic units, as measured by accumulation of neutral mutations. In a recent evolutionary reconstruction, all extant *S. aureus* clonal complexes tested had inferred last common ancestors in the past 250 years, most much sooner^49^, suggesting frequent turnover of new strains. The reasons for these replacement events could be a unique historical feature of the past 2-3 centuries, caused perhaps by the development of human healthcare systems and the changing chemical environment of human and animal microhabitats due to technological advances but the pattern of frequent strain replacement seems common to many bacterial species^19^. Possibly, strains are replaced from within by the wavelike expansion of successful clones. Something like this process may be happening with the expansion of USA300 since the late 1980s, gradually becoming the most common CC8 strain in the USA ^62,63^. This explanation implies that strains occupy distinct niches, with adaptation possibly defined by the composition of their non-core genes ^64,65^. Substrains would then be competing with each other to occupy the strain niche. There is not strong evidence of distinct within-host niches for most *S. aureus* strains but there are clear associations of strains with particular animal hosts^66^. New strains can also emerge from outside by genome-scale recombination events, exemplified by CC239 strains, which were formed by recombination of a large segment of a CC30 chromosome into a CC8 background ^22,23^. Judging by the relatively small size of the “99.1-99.5% bump” (**Figure 1**) these types of events may be a rare but ongoing process.

The second question we highlight concerns the functional implications of the partition of strain-concentrated and strain-diffuse genes. There is a bias for deletion in bacterial genomes^67^ that implies genes maintained over time are under enduring strong selection. Conversely, the strain-diffuse gene pattern can be seen as cycles of gene gain under neutral selection (i.e. driven by gene transfer alone) or short term positive selection followed by rapid removal. However we do not know of any studies that address the underlying reasons for the difference in strain-level versus substrain-level selection. Toxins are interesting in this regard because of their importance for *S. aureus* virulence. Why are some toxins maintained as core functions (e.g alpha-toxin (*hly)*), some strain-concentrated (e.g enterotoxin G (seg)) and some strain-diffuse, present in diverse substrains (e.g Panton-Valentine leukocidin (*lukFS)*)? (**Figure 4**). The superantigen-type toxins are split between strain-concentrated and strain-diffuse genes, suggesting that former functions may be strongly linked to strain niches. Related to these issues is the question of long-term maintenance of diversity of strain-concentrated genes under conditions of relatively low transfer rate and rapid strain extinction that would suggest a high rate of stochastic loss. Could there be frequency-dependent selection operating across the *S. aureus* species on strain-concentrated genes?.

In summary, this work revealed a new partition in the structure of the *S. aureus* pangenome that will spur further studies on genome evolution and subspeciation in the species. The methodology for refining large amounts of public data, defining strains using ANI and following strain-specificity of the pangenome using F_st_ can also be applied to other bacterial species. Comparisons to other species, particularly from the *Staphylococcus* genus, will reveal the commonalities and unique selective pressures acting on the pangenome of this dangerous pathogen.

## Acknowledgements

D.B.W. and T.D.R. were supported by an Emory University Synergy II_Nexus / MP3 award. T.D.R. was supported by funding from NIH awards AI158452 and AI139188. V.R. was supported by NIH AI139188 and the NIH T32 AI138952 award “Infectious Disease Across Scales Training Program (IDASTP)”. D.B.W was supported by funding from the Simons Foundation (Mathematical Modeling of Living Systems Investigator award 508600), the Sloan Foundation (Research Fellowship FG-2021-16667), and the NSF (award 2146260). We would like to thank Megan Phillips and Anayancy Ramos Facio for discussions about the manuscript.

## Methods

### Public genome collection, processing and filtering

Bactopia v1.7.0 was used to download and process all genomes used in this dataset. Bactopia is a software pipeline for comprehensive analysis of bacterial genomes based on Nextflow ^68,69^. The command “*bactopia search “Staphylococcus aureus” --prefix saureus”* was used to download all *S. aureus* short-read sequences available on Sequence Read Archive (SRA) as of September 2022. Bactopia used SKESA to assemble genomes, Bakta to annotate and Snippy for variant calling ^70,71^. Assembly quality was evaluated using QUAST and CheckM ^72,73^. *S. aureus* CC and ST were based on the pubmlst database ^20^. (https://pubmlst.org/bigsdb?db=pubmlst_saureus_seqdef&page=downloadProfiles&scheme_id=1). AgrVATE v1.0.5 was used to assign *agr* types ^15^. Only samples having greater than 50× coverage, mean per-read quality greater than 20, mean read length greater than 75 bp, and an assembly with less than 200 contigs were considered for the analysis (corresponding to ‘Gold’ and ‘Silver’ ranks as designated by Bactopia. Samples that were detected as not *S. aureus* according to kmer based identification or CheckM were then removed. Coverage for all samples were capped at 100x. For every sample, bactopia performs variant calling using Snippy against an auto-chosen reference sequence based on the smallest MASH distance to a complete *S. aureus* genome in RefSeq ^70,74^. For each variant identified, the allele frequencies were calculated from the bam files using bcftools mpileup ^75^. Samples having average minor allele frequency > 0.05 were considered mixed strains and therefore removed. Samples having total number of variants > 150,000 compared to the auto-chosen reference (or more than 5% of the genome) were also considered non-*S. aureus* and removed ^76^. This process reduced 83,383 samples to 56,771. Since Bactopia collected and processed only short read *S. aureus* data, we added complete *S. aureus* genome sequences to this set. Out of 1,475 complete genomes publicly available as of February 2023, 1,263 did not have any ‘N’ characters in their assemblies and were added to the filtered dataset of 56,771, leading to a total of 58,034 genomes. The 212 complete genomes containing ‘N’ characters were not used in this study.

### Substrain dereplication

Samples were grouped by their MLST types as assigned by Bactopia and for each ST, an all vs all MASH distance estimation ^74^ was run. Samples with a MASH distance < 0.0005 were grouped into clusters and a random genome was chosen as the cluster representative ^16^. However, where possible, we used complete genomes as the cluster representative. Samples with unassigned STs were grouped together and treated the same. The resulting final dereplicated set comprised 7954 genomes and was used for pangenome construction.

### Pangenome analysis

The bakta annotation produced by the original Bactopia run was used as input for pangenome estimation with PIRATE 1.0.5 ^17^. PIRATE was run using default parameters with the additional flags -a to obtain core genome alignments and -k “--diamond” to use DIAMOND for the amino-acid sequence comparisons ^77^. SNP-sites v 2.5.1 ^78^ was run on the PIRATE core genome alignment to extract only polymorphic sites (709,911 sites) and the resulting alignment was used to construct a core genome phylogeny with FastTree v 2.1.11 ^79^(GTR model, 1000 bootstrap resamples). The phylogeny was visualized using the R package ggtree ^80,81^. We used Homoplasyfinder^27^ to count the number of state changes of each non-core gene on the phylogeny. geNomad v1.5^28^ was used to predict mobile genetic elements.

### Strain definition based on ANI

All-vs-All pairwise ANI was calculated for the 7,954 dereplicated genomes using fastANI v1.33 ^76^. Strain assignments were performed based on average linkage hierarchical clustering and samples that had ANI 99.5% or greater were clustered together. The average ANI of each genome with every other genome in a given cluster was calculated and the genome with the highest average ANI was assigned as the strain representative.

### Calculating F_ST_

We created a custom R function to calculate the F_ST_ for each gene, with group membership defined as strain type, clonal complex or *agr* group, depending on the purpose of the comparison. The input was a binary presence/absence data frame, with genes as columns and genomes as rows. F_ST_ was calculated using Weir’s formula ^24^.

### Creating the 740-set and 740-set-90 pangenomes

We randomly subsampled the 37 strains with > 20 substrains to 20 substrain genomes each. We rerun PIRATE 1.0.5 with default parameters and created a core pangnome tree using FastTree v 2.1.11 as described above. To create the “740-set-90” pangenome we the 740 genomes through PIRATE 1.0.5 with minimum clustering threshold of 90% amino acid identity.

### Chromosomal locations of non-core genes

We used two methods for mapping chromosomal locations of non-core genes based on the co-ords output of the PIRATE 1.0.5 pipeline for the 7954-set and 740-set pangenomes. First we screened 377 complete substrain genome that had *dnaA* as their first gene by BLAST and collated the start coordinate of each non-core gene. The second method was to collate the start coordinate of nearest core gene on the same contig as each non-core gene. For each class of non-core gene 20,000 random genes were selected as well as a control of 20,000 genes of all classes (including core). If the non-core gene was on a contig that did not have a core gene then its status was returned as “unlinked”.

### Antibiotic resistance, virulence and phage defense functions

To assign antibiotic-resistance genes we queried representative protein sequences of each gene family of the 7954-set produced by PIRATE against the AMRFinder+^31^ database using tblastn ^82^ with a threshold of >= 90% identity as a match. We filtered the out virulence-associated genes using matches the terms: “serine_protease”, “enterotoxin”, “hemolysin”, “Panton”, “adhesin”, “complement”, “aureolysin”, “exfoliative”, “toxin”, “intracellular_survival”, “serum_survival” and “leukocidin” and the kept the remainder as antibiotic-resistance gene matches. To assign phage defense related functions, we queried the 7954-set representative proteins against the online defensefinder database^33^ (https://defense-finder.mdmparis-lab.com/) on 2023-10-17.

### Statistical analysis and data visualization

All statistics and tSNE were performed in R using package rstatix ^83^. All plots were visualized using R package ggplot2 ^84^. Other visualizations were performed using draw.io and Sakneymatic ^85,86^.

## Data availability

PIRATE pangenome outputs, genes and strain lists and representative genome sets are available on Zenodo https://zenodo.org/records/10471309.

## Supplemental Data

**Figure S1:**
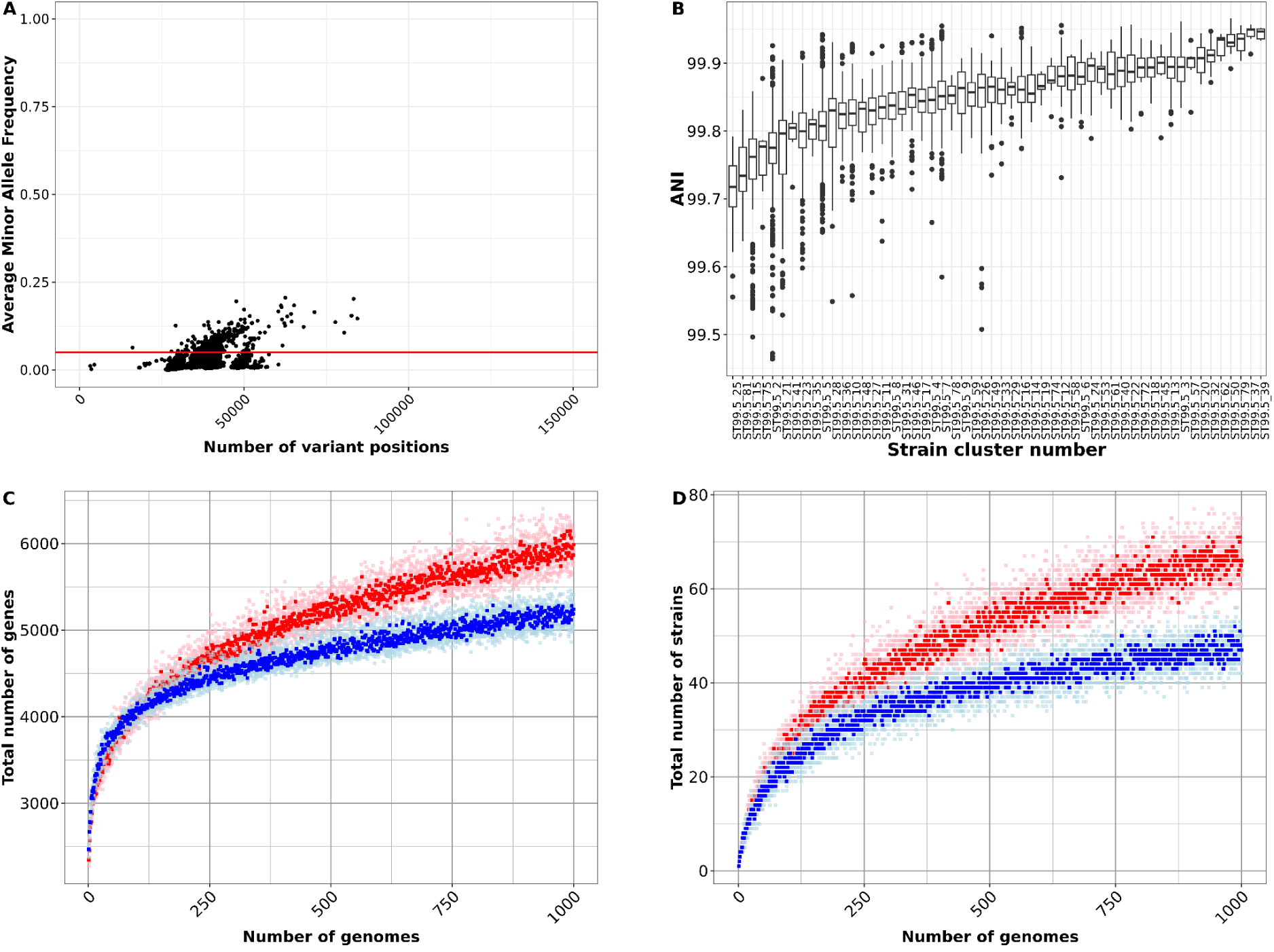
Effect of filtering, clustering and dereplicating 83,383 *S. aureus* genomes. (A) The x-axis shows the total number of variants when compared with the Bactopia auto-chosen reference, and the y-axis shows the average minor allele frequency (MAF). Each dot is one of 57,093 genomes which were obtained after filtering out samples ranked ‘Bronze’ or ‘Exclude’ by Bactopia and/or found to have non-*S. aureus* genome content by Bactopia and CheckM (**Figure 1**). Samples in the top quadrant (Above red horizontal line - Average MAF > 0.05) were considered to be *S. aureus* strain mixtures and were discarded. The remaining 56,771 samples in the bottom quadrant (< 0.05 Average MAF) were used for further analysis. (B) Boxplots showing spread of pairwise ANI within each “strain” cluster. Only strain clusters having more than 10 genomes are shown. Black horizontal line within each boxplot shows the median within strain-cluster pairwise ANI. Total number of unique genes discovered (C) and total number of strains discovered (D) for every new genome added from the dereplicated set (red dots) or a random genome from the un-dereplicated 58,034 (blue dots). Up to 1000 random genomes were added from each set and the total number of unique genes or strains were measured for every genome added (light red and light blue dots). This procedure was repeated 5 times and the median number of genes or strains discovered are shown in dark red and dark blue dots. More genes and more strains were discovered from the same number of genomes (after observing 1000 genomes) in the dereplicated set compared to the un-dereplicated set.

**Figure S2:**
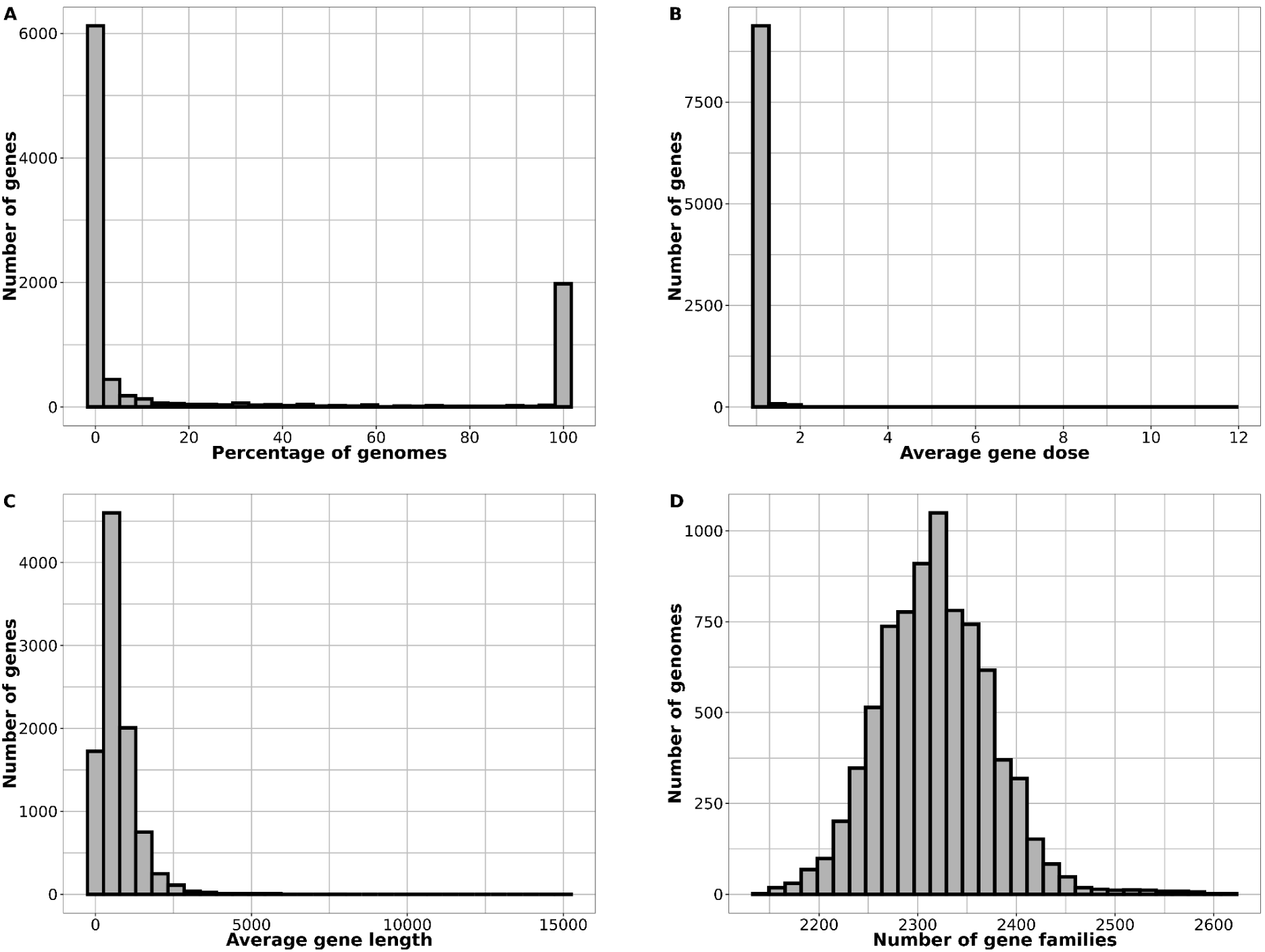
The 7,954 substrain pangenome of *S. aureus*. Histograms depicting the (A) frequency distribution of genes in our dataset, (B) the average dosage of each gene per genome, (C) the average length distribution of each gene, and (D) the distribution of the number of unique PIRATE gene families per genome.

**Figure S3:**
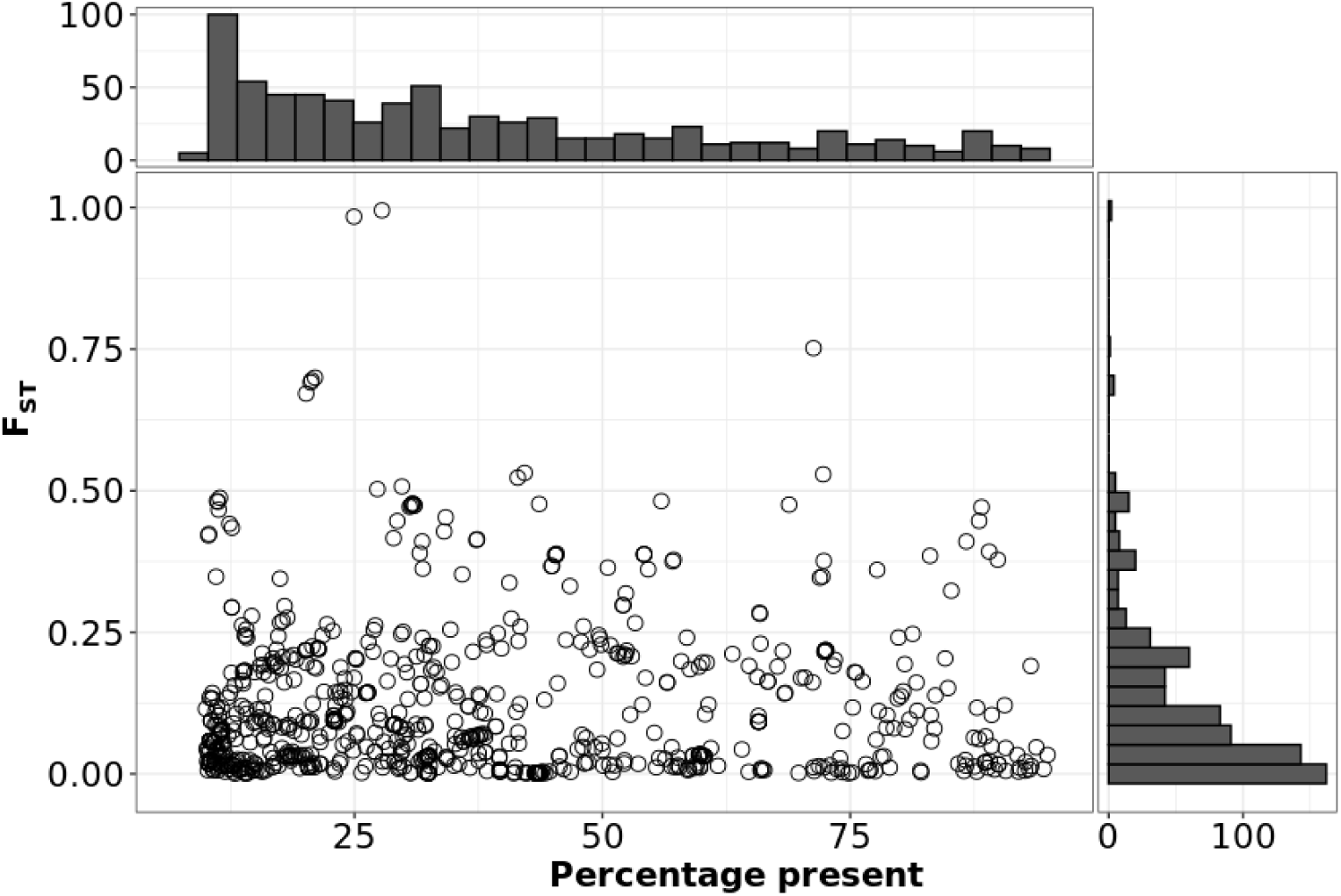
There are no *agr* group specific intermediate genes aside from *agrD*. Dot plot showing percentage prevalence of only intermediate genes (> 10%, < 95%) on the x-axis and the corresponding F_ST_ on the y-axis. F_ST_ scores calculated for *agr* type-based population segregation. The three dots > 0.75 F_ST_ correspond to the *agrD*, which are known to be lineage specific. The *agrD* of the fourth *agr* type is absent in this plot as it is present in < 10% of the population 15.

**Figure S4:**
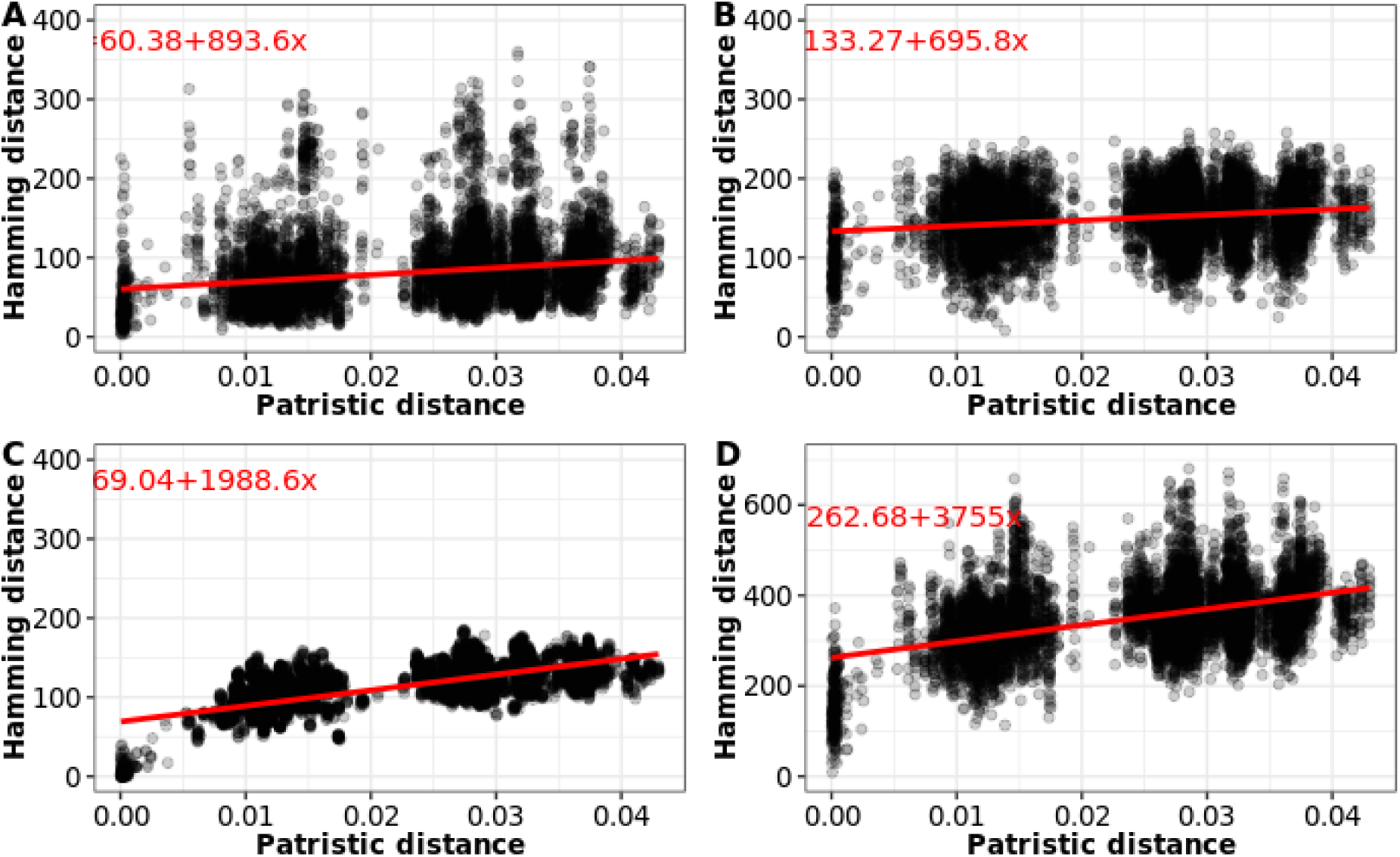
strain-concentrated gene content declines gradually with core-gene distance. Each dot represents a comparison between substrains in the 740-set. Patristic distance was tip-tip distance on the phylogeny. Hamming distance was calculated from a presence absence matrix of each non-core gene type: A) rare genes, B) strain-diffuse, C) strain-concentrated, D) all non-core (note different y-axis scale). Red lines show the linear model fit.

**Figure S5:**
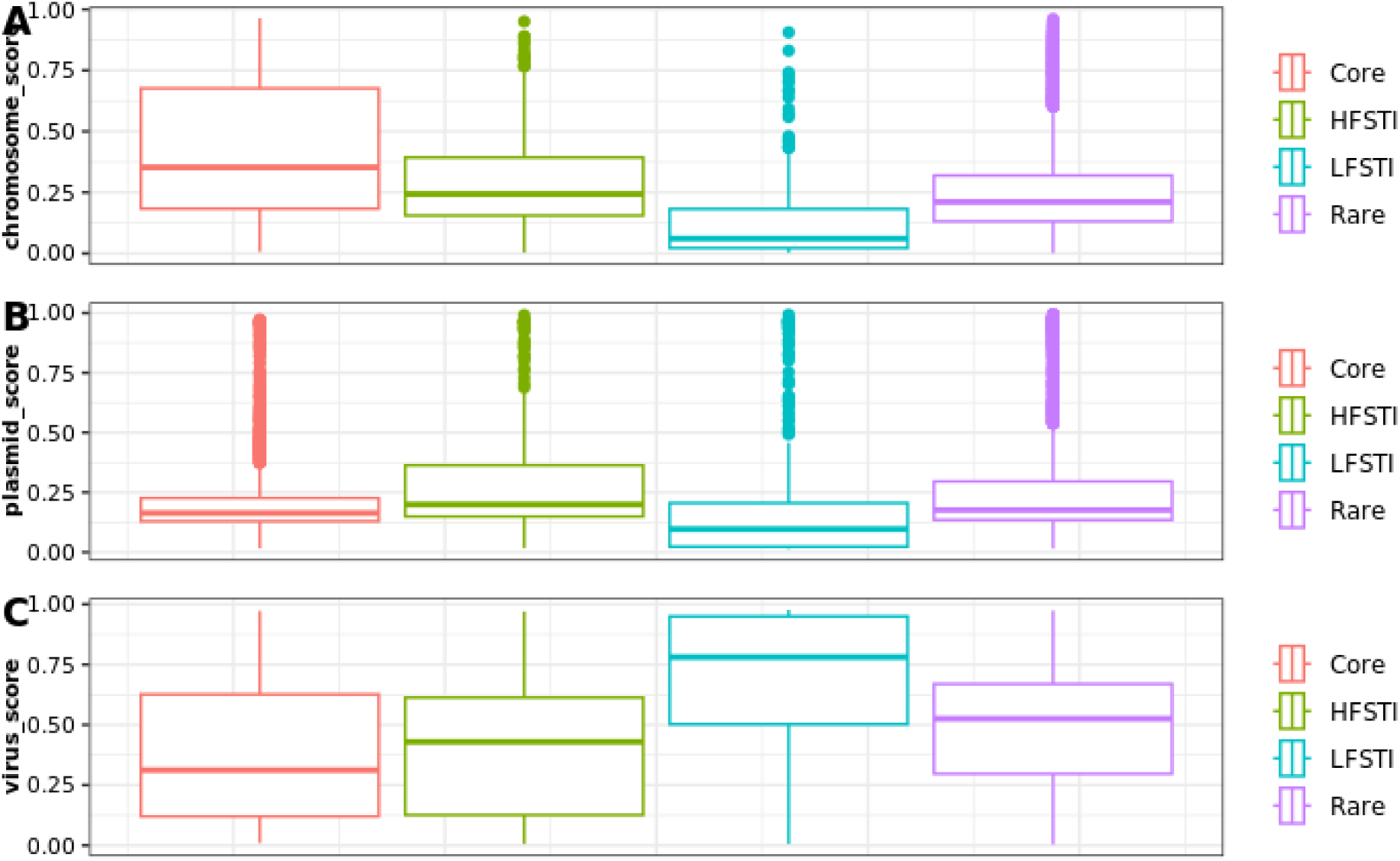
Genomad score distributions for 7954-set pangenomes. The geNomad ^28^ probability scores for A) chromosome B) plasmid and C) virus were grouped by gene class. All differences were significant in a Tukey’s pairwise comparisons at < 0.05 (corrected for multiple tests), except strain-diffuse-Core plasmid_score and strain-concentrated-Core virus score.

**Figure S6.**
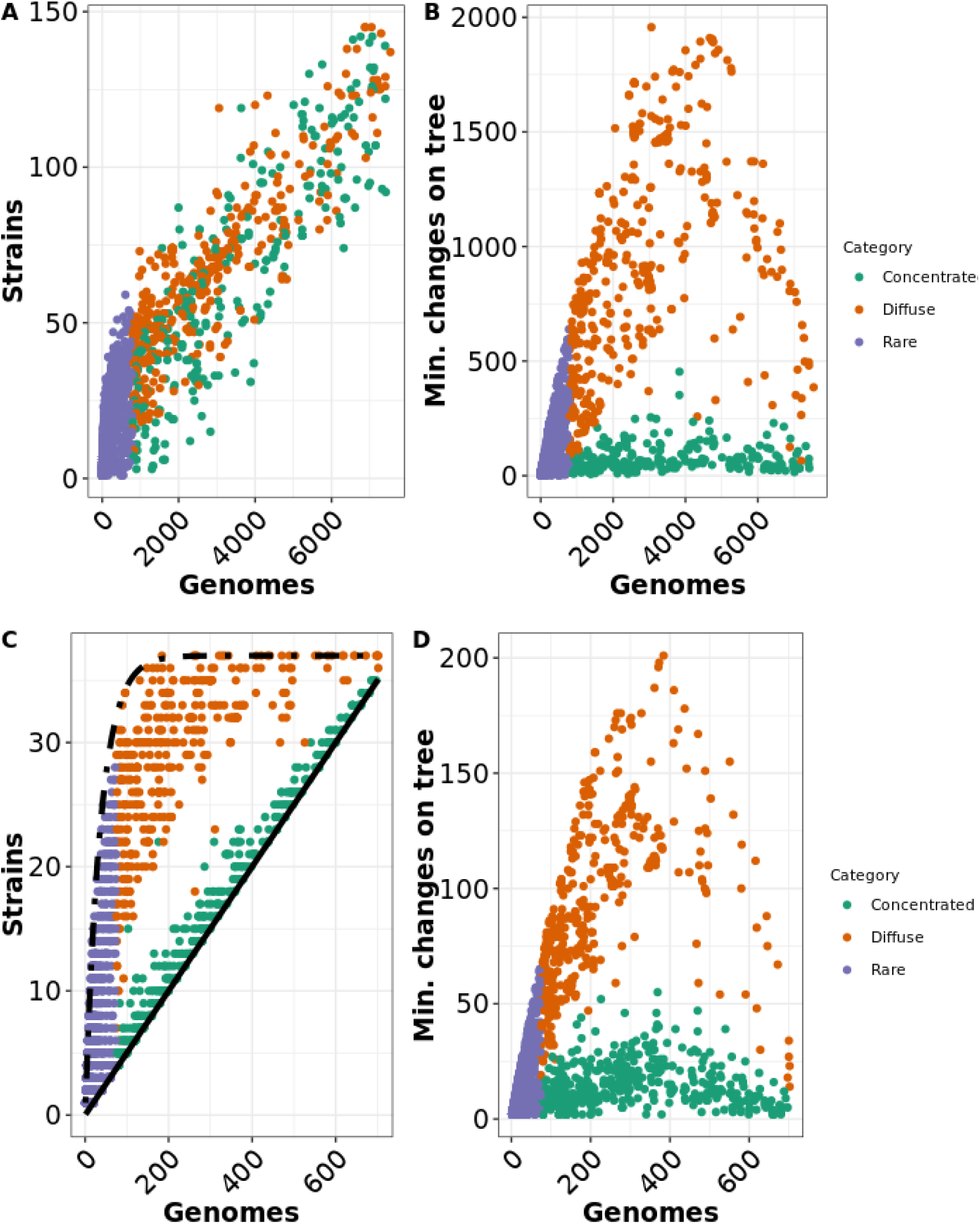
Relationship between gene prevalence, number of strains and homoplasy for non-core genes for the 7954-set (a,b) and the 740-set-90 (c,d) The plots are formatted as **Figure 5**. Each dot represents a non-core gene. “concentrated” = strain-concentrated, “diffuse” = strain-diffuse. Panels A and C show the relationship between overall prevalence (number of genomes out of 740) and number of strains (out of 37) each gene is found in. Panels B and D show the relationship between prevalence of estimated number of changes on the species tree calculated by homoplasyfinder^27^. In panel A, the unbalanced nature of the 7954-set (a few strains have thousands of genomes, many have only one) obscures the differences between concentrated and diffuse: it not possible to plot simple bounds of lowest possible and random gene distribution into strains as it is for the 740-90 set (panel C).

**Figure S7:**
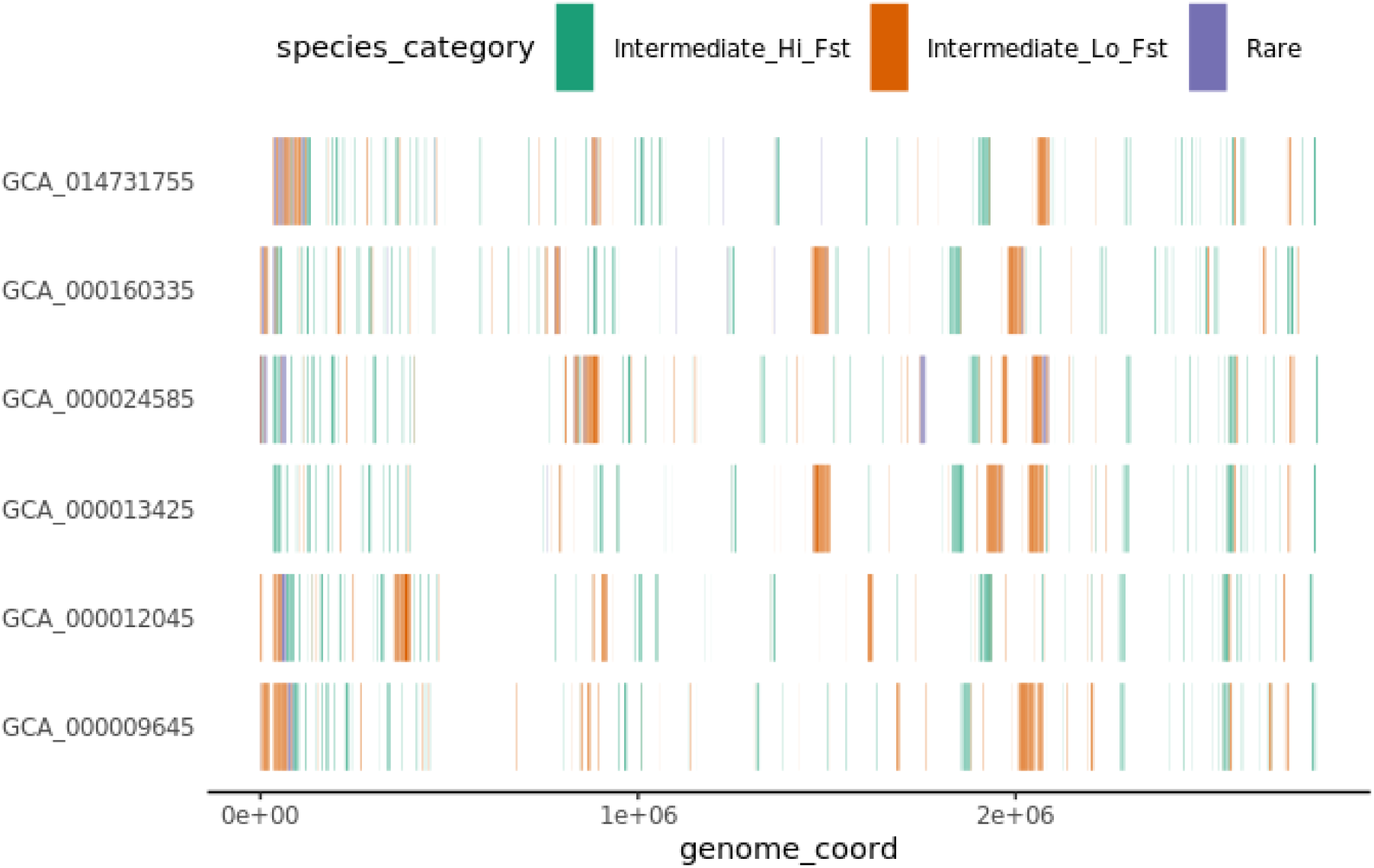
Chromosome start locations of non-core genes on six *S. aureus* complete chromosomes. The name on the left-hand side refers to NCBI assembly database designations. GCA_014731755 is CC30 MRSA; GCA_000160335 is CC30 MSSA; GCA_000024585 is CC5 MSSA; GCA_0000134525 is CC8 MRSA; GCA_000012045 is CC8 MRSA; GCA_000009645 is CC5 MRSA (N315 the S. aureus type strain). “Intermediate_Hi_Fst” = strain-concentrated; “Intermediate_Lo_Fst” = strain-diffuse.

**Table S1:** *S. aureus* studies quoting pangenome statistics. “?” indicates that the corresponding information could not be found

| Title | Date | No. of genomes | Sampling space | Assembly level | Pangenome tool | No. core | Total gene families |
| --- | --- | --- | --- | --- | --- | --- | --- |
| Comparative Pan-Genomic Analysis Revealed an Improved Multi-Locus Sequence Typing Scheme for *Staphylococcus aureus* <sup>87</sup> | 2022-11-19 | 502 | Diverse | Complete | PanRV (Roary) | 2320 | 12477 |
| Pan-Genome Analysis of *Staphylococcus aureus* Reveals Key Factors Influencing Genomic Plasticity <sup>88</sup> | 2022-11-01 | 1519 | Diverse | All | Roary | 1000 | 16794 |
| Pangenomic Approach To Understanding Microbial Adaptations within a Model Built Environment, the International Space Station, Relative to Human Hosts and Soil <sup>50</sup> | 2022-01-08 | 106 | ISS, human, soil | All | Roary | 1935 | 6847 |
| The Epidemiological and Pangenome Landscape of *Staphylococcus aureus* and Identification of Conserved Novel Candidate Vaccine Antigens <sup>47</sup> | 2022-02-01 | 355 | Diverse | All | ? | 2025 | 7199 |
| Analyses of Livestock-Associated *Staphylococcus aureus* Pan-Genomes Suggest Virulence Is Not Primary Interest in Evolution of Its Genome <sup>51</sup> | 2019-05-22 | 14 | Livestock associated | Complete | Roary | 1969 | 4637 |
| Comparative genome-scale modelling of *Staphylococcus aureus* strains identifies strain-specific metabolic capabilities linked to pathogenicity <sup>89</sup> | 2016-06-10 | 64 | Diverse | All | dGenome DuctAPE | 1441 | 7457 |
| PanRV: Pangenome-reverse vaccinology approach for identifications of potential vaccine candidates in microbial pangenome <sup>48</sup> | 2019-03-12 | 301 | Diverse | All | PanRV (Roary) | 1524 | 11384 |
| Whole-Genome Sequencing of *Staphylococcus aureus* and *Staphylococcus haemolyticus* Clinical Isolates from Egypt <sup>43</sup> | 2022-06-21 | 90 | 56 from 1 hospital and 34 from greater Arab region | All | Anvio | 1501 | 4283 |
| Phylogenomic Analysis Reveals the Evolutionary Route of Resistant Genes in *Staphylococcus aureus* <sup>52</sup> | 2019-11-03 | 152 | Diverse | Complete | Manual alignment and clustering | 2426 | 6326 |
| Comparative genomic analysis of *Staphylococcus aureus* isolates associated with either bovine intramammary infections or human infections demonstrates the importance of restriction-modification systems in host adaptation <sup>90</sup> | 2022-02-18 | 187 | Human and cattle | All | Roary | 2700 | 6812 |
| Molecular Epidemiology of Staphylococcus aureus in China Reveals the Key Gene Features Involved in Epidemic Transmission and Adaptive Evolution <sup>44</sup> | 2022-10-03 | 332 | Human clinical strains from China | All | Heap's law algorithms | 890 | 5832 |
| Estimated Roles of the Carrier and the Bacterial Strain When Methicillin-Resistant Staphylococcus aureus Decolonization Fails: a Case-Control Study <sup>45</sup> | 2022-08-24 | 477 | MRSA carriers from Denmark hospitals | All | panX | 1671 | 5925 |
| Forecasting Staphylococcus aureus Infections Using Genome-Wide Association Studies, Machine Learning, and Transcriptomic Approaches <sup>91</sup> | 2022-07-05 | 356 | Mostly human | All | Panaroo | 1489 | 8827 |
| Carriage prevalence and genomic epidemiology of Staphylococcus aureus among Native American children and adults in the Southwestern USA <sup>46</sup> | 2022-05-13 | 92 | Native Americans from Southwestern USA | Complete | Roary | 1808 | ? |
| Polyclonality, Shared Strains, and Convergent Evolution in Chronic Cystic Fibrosis Staphylococcus aureus Airway Infection <sup>42</sup> | 2020-03-23 | 1382 | Longitudinal sampling from 246 children with CF from the US | All | Roary | 1142 | 21358 |
| PIRATE: A fast and scalable pangenomics toolbox for clustering diverged orthologues in bacteria <sup>17</sup> | 2019-10-09 | 253 | Diverse | All | PIRATE | 2433 | 4250 |
| Whole-Genome Sequencing for Routine Pathogen Surveillance in Public Health: a Population Snapshot of Invasive Staphylococcus aureus in Europe <sup>92</sup> | 2016-05-05 | 308 | Invasive isolates from Europe hospitals within a 6 month period | All | BlastP & TribeMCL | ? | 4281 |

## References

1 Kourtis AP, Hatfield K, Baggs J, Mu Y, See I, Epson E, et al. Vital Signs: Epidemiology and Recent Trends in Methicillin-Resistant and in Methicillin-Susceptible Staphylococcus aureus Bloodstream Infections - United States. MMWR Morb Mortal Wkly Rep 2019;68:214–9. 10.15585/mmwr.mm6809e1.

2 Tettelin H, Masignani V, Cieslewicz MJ, Donati C, Medini D, Ward NL, et al. Genome analysis of multiple pathogenic isolates of Streptococcus agalactiae: implications for the microbial ‘pan-genome’. Proc Natl Acad Sci U S A 2005;102:13950–5. 10.1073/pnas.0506758102.

3 Howden BP, Giulieri SG, Wong Fok Lung T, Baines SL, Sharkey LK, Lee JYH, et al. Staphylococcus aureus host interactions and adaptation. Nat Rev Microbiol 2023. 10.1038/s41579-023-00852-y.

4 Vestergaard M, Frees D, Ingmer H. Antibiotic Resistance and the MRSA Problem. Microbiol Spectr 2019;7.: 10.1128/microbiolspec.GPP3-0057-2018.

5 Peschel A, Otto M. Phenol-soluble modulins and staphylococcal infection. Nat Rev Microbiol 2013;11:667–73. 10.1038/nrmicro3110.

6 Spaan AN, van Strijp JAG, Torres VJ. Leukocidins: staphylococcal bi-component pore-forming toxins find their receptors. Nat Rev Microbiol 2017. 10.1038/nrmicro.2017.27.

7 Azarian T, Cella E, Baines SL, Shumaker MJ, Samel C, Jubair M, et al. Genomic Epidemiology and Global Population Structure of Exfoliative Toxin A-Producing Staphylococcus aureus Strains Associated With Staphylococcal Scalded Skin Syndrome. Front Microbiol 2021;12:2307. 10.3389/fmicb.2021.663831.

8 Spoor LE, McAdam PR, Weinert LA, Rambaut A, Hasman H, Aarestrup FM, et al. Livestock origin for a human pandemic clone of community-associated methicillin-resistant Staphylococcus aureus. MBio 2013;4.: 10.1128/mBio.00356-13.

9 Su M, Lyles JT, Petit RA Iii, Peterson J, Hargita M, Tang H, et al. Genomic analysis of variability in Delta-toxin levels between Staphylococcus aureus strains. PeerJ 2020;8:e8717. 10.7717/peerj.8717.

10 Benson MA, Ohneck EA, Ryan C, Alonzo F 3rd, Smith H, Narechania A, et al. Evolution of hypervirulence by a MRSA clone through acquisition of a transposable element. Mol Microbiol 2014;93:664–81. 10.1111/mmi.12682.

11 Su M, Davis MH, Peterson J, Solis-Lemus C, Satola SW, Read TD. Effect of genetic background on the evolution of Vancomycin-Intermediate Staphylococcus aureus (VISA). PeerJ 2021;9:e11764. 10.7717/peerj.11764.

12 Petit RA 3rd, Read TD. Staphylococcus aureus viewed from the perspective of 40,000+ genomes. PeerJ 2018;6:e5261. 10.7717/peerj.5261.

13 Van Rossum T, Ferretti P, Maistrenko OM, Bork P. Diversity within species: interpreting strains in microbiomes. Nat Rev Microbiol 2020. 10.1038/s41579-020-0368-1.

14 Liao H, Ji Y, Sun Y. High-resolution strain-level microbiome composition analysis from short reads. Microbiome 2023;11:183. 10.1186/s40168-023-01615-w.

15 Raghuram V, Alexander AM, Loo HQ, Petit RA 3rd, Goldberg JB, Read TD. Species-Wide Phylogenomics of the Staphylococcus aureus Agr Operon Revealed Convergent Evolution of Frameshift Mutations. Microbiol Spectr 2022:e0133421. 10.1128/spectrum.01334-21.

16 Raghuram V, Read T. Help, I have too many genome sequences!. 2022. 10.5281/zenodo.7278310.

17 Bayliss SC, Thorpe HA, Coyle NM, Sheppard SK, Feil EJ. PIRATE: A fast and scalable pangenomics toolbox for clustering diverged orthologues in bacteria. Gigascience 2019;8:598391. 10.1093/gigascience/giz119.

18 Planet PJ, Narechania A, Chen L, Mathema B, Boundy S, Archer G, et al. Architecture of a Species: Phylogenomics of Staphylococcus aureus. Trends Microbiol 2016. 10.1016/j.tim.2016.09.009.

19 Rodriguez-R LM, Conrad RE, Viver T, Feistel DJ, Lindner BG, Venter SN, et al. An ANI gap within bacterial species that advances the definitions of intra-species units. MBio 2023:e0269623. 10.1128/mbio.02696-23.

20 Jolley KA, Bray JE, Maiden MCJ. Open-access bacterial population genomics: BIGSdb software, the PubMLST.org website and their applications. Wellcome Open Res 2018;3:124. 10.12688/wellcomeopenres.14826.1.

21 Yousuf B, Flint A, Weedmark K, McDonald C, Bearne J, Pagotto F, et al. Genome Sequence of Staphylococcus aureus Strain PS/BAC/317/16/W, Isolated from Contaminated Platelet Concentrates in England. Microbiol Resour Announc 2021;10:e0057721. 10.1128/MRA.00577-21.

22 Gill JL, Hedge J, Wilson DJ, MacLean RC. Evolutionary Processes Driving the Rise and Fall of Staphylococcus aureus ST239, a Dominant Hybrid Pathogen. MBio 2021;12:e0216821. 10.1128/mBio.02168-21.

23 Robinson DA, Enright MC. Evolution of Staphylococcus aureus by large chromosomal replacements. J Bacteriol 2004;186:1060–4.

24 Weir BS. Estimating F-statistics: A historical view. Philos Sci 2012;79:637–43. 10.1086/667904.

25 Melles DC, van Leeuwen WB, Boelens HAM, Peeters JK, Verbrugh HA, van Belkum A. Panton-Valentine leukocidin genes in Staphylococcus aureus. Emerg Infect Dis 2006;12:1174–5. 10.3201/eid1207.050865.

26 Krakauer T. Staphylococcal Superantigens: Pyrogenic Toxins Induce Toxic Shock. Toxins 2019;11.: 10.3390/toxins11030178.

27 Crispell J, Balaz D, Gordon SV. HomoplasyFinder: a simple tool to identify homoplasies on a phylogeny. Microb Genom 2019;5.: 10.1099/mgen.0.000245.

28 Camargo AP, Roux S, Schulz F, Babinski M, Xu Y, Hu B, et al. Identification of mobile genetic elements with geNomad. Nat Biotechnol 2023. 10.1038/s41587-023-01953-y.

29 Gill SR, Fouts DE, Archer GL, Mongodin EF, Deboy RT, Ravel J, et al. Insights on evolution of virulence and resistance from the complete genome analysis of an early methicillin-resistant Staphylococcus aureus strain and a biofilm-producing methicillin-resistant Staphylococcus epidermidis strain. J Bacteriol 2005;187:2426–38. 10.1128/JB.187.7.2426-2438.2005.

30 Warne B, Harkins CP, Harris SR, Vatsiou A, Stanley-Wall N, Parkhill J, et al. The Ess/Type VII secretion system of Staphylococcus aureus shows unexpected genetic diversity. BMC Genomics 2016;17:222. 10.1186/s12864-016-2426-7.

31 Feldgarden M, Brover V, Gonzalez-Escalona N, Frye JG, Haendiges J, Haft DH, et al. AMRFinderPlus and the Reference Gene Catalog facilitate examination of the genomic links among antimicrobial resistance, stress response, and virulence. Sci Rep 2021;11:12728. 10.1038/s41598-021-91456-0.

32 Chambers HF, Deleo FR. Waves of resistance: Staphylococcus aureus in the antibiotic era. Nat Rev Microbiol 2009;7:629–41. 10.1038/nrmicro2200.

33 Tesson F, Hervé A, Mordret E, Touchon M, d’Humières C, Cury J, et al. Systematic and quantitative view of the antiviral arsenal of prokaryotes. Nat Commun 2022;13:2561. 10.1038/s41467-022-30269-9.

34 Xia G, Wolz C. Phages of Staphylococcus aureus and their impact on host evolution. Infect Genet Evol 2014;21:593–601. 10.1016/j.meegid.2013.04.022.

35 McCarthy AJ, Lindsay JA. The distribution of plasmids that carry virulence and resistance genes in Staphylococcus aureus is lineage associated. BMC Microbiol 2012;12:104. 10.1186/1471-2180-12-104.

36 Beghini F, McIver LJ, Blanco-Miguez A, Dubois L, Asnicar F, Maharjan S, et al. Integrating taxonomic, functional, and strain-level profiling of diverse microbial communities with bioBakery 3. Cold Spring Harbor Laboratory 2020:2020.11.19.388223. 10.1101/2020.11.19.388223.

37 Jin X, Yu FB, Yan J, Weakley AM, Dubinkina V, Meng X, et al. Culturing of a complex gut microbial community in mucin-hydrogel carriers reveals strain- and gene-associated spatial organization. Nat Commun 2023;14:3510. 10.1038/s41467-023-39121-0.

38 Maiden MC, Bygraves JA, Feil E, Morelli G, Russell JE, Urwin R, et al. Multilocus sequence typing: a portable approach to the identification of clones within populations of pathogenic microorganisms. Proc Natl Acad Sci U S A 1998;95:3140–5. 10.1073/pnas.95.6.3140.

39 Maiden MCJ, van Rensburg MJJ, Bray JE, Earle SG, Ford SA, Jolley KA, et al. MLST revisited: the gene-by-gene approach to bacterial genomics. Nat Rev Microbiol 2013;11:728–36. 10.1038/nrmicro3093.

40 Falush D. Toward the use of genomics to study microevolutionary change in bacteria. PLoS Genet 2009;5:e1000627. 10.1371/journal.pgen.1000627.

41 Jamrozy DM, Harris SR, Mohamed N, Peacock SJ, Tan CY, Parkhill J, et al. Pan-genomic perspective on the evolution of the Staphylococcus aureus USA300 epidemic. Microb Genom 2016;2:e000058. 10.1099/mgen.0.000058.

42 Long DR, Wolter DJ, Lee M, Precit M, McLean K, Holmes E, et al. Polyclonality, Shared Strains, and Convergent Evolution in Chronic CF S. aureus Airway Infection. Am J Respir Crit Care Med 2020. 10.1164/rccm.202003-0735OC.

43 Montelongo C, Mores CR, Putonti C, Wolfe AJ, Abouelfetouh A. Whole-Genome Sequencing of Staphylococcus aureus and Staphylococcus haemolyticus Clinical Isolates from Egypt. Microbiol Spectr 2022;10:e0241321. 10.1128/spectrum.02413-21.

44 Xu Z, Yuan C. Molecular Epidemiology of Staphylococcus aureus in China Reveals the Key Gene Features Involved in Epidemic Transmission and Adaptive Evolution. Microbiol Spectr 2022:e0156422. 10.1128/spectrum.01564-22.

45 Holm MKA, Jørgensen KM, Bagge K, Worning P, Pedersen M, Westh H, et al. Estimated Roles of the Carrier and the Bacterial Strain When Methicillin-Resistant Staphylococcus aureus Decolonization Fails: a Case-Control Study. Microbiol Spectr 2022:e0129622. 10.1128/spectrum.01296-22.

46 Cella E, Sutcliffe CG, Tso C, Paul E, Ritchie N, Colelay J, et al. Carriage prevalence and genomic epidemiology of Staphylococcus aureus among Native American children and adults in the Southwestern USA. Microbial Genomics 2022;8:000806. 10.1099/mgen.0.000806.

47 Naz K, Ullah N, Naz A, Irum S, Dar HA, Zaheer T, et al. The Epidemiological and Pangenome Landscape of Staphylococcus aureus and Identification of Conserved Novel Candidate Vaccine Antigens. Curr Proteomics 2022;19:114–26. 10.2174/1570164618666210212122847.

48 Naz K, Naz A, Ashraf ST, Rizwan M, Ahmad J, Baumbach J, et al. PanRV: Pangenome-reverse vaccinology approach for identifications of potential vaccine candidates in microbial pangenome. BMC Bioinformatics 2019;20:123. 10.1186/s12859-019-2713-9.

49 Yebra G, Harling-Lee JD, Lycett S, Aarestrup FM, Larsen G, Cavaco LM, et al. Multiclonal human origin and global expansion of an endemic bacterial pathogen of livestock. Proc Natl Acad Sci U S A 2022;119:e2211217119. 10.1073/pnas.2211217119.

50 Blaustein RA, McFarland AG, Ben Maamar S, Lopez A, Castro-Wallace S, Hartmann EM. Pangenomic Approach To Understanding Microbial Adaptations within a Model Built Environment, the International Space Station, Relative to Human Hosts and Soil. mSystems 2019;4:e00281–18. 10.1128/mSystems.00281-18.

51 Rao RT, Sivakumar N, Jayakumar K. Analyses of Livestock-Associated Staphylococcus aureus Pan-Genomes Suggest Virulence Is Not Primary Interest in Evolution of Its Genome. OMICS 2019;23:224–36. 10.1089/omi.2019.0005.

52 John J, George S, Nori SRC, Nelson-Sathi S. Evolutionary route of resistant genes in Staphylococcus aureus. Genome Biol Evol 2019. 10.1093/gbe/evz213.

53 Vernikos G, Medini D, Riley DR, Tettelin H. Ten years of pan-genome analyses. Curr Opin Microbiol 2014;23C:148–54. 10.1016/j.mib.2014.11.016.

54 Schwengers O, Jelonek L, Dieckmann MA, Beyvers S, Blom J, Goesmann A. Bakta: rapid and standardized annotation of bacterial genomes via alignment-free sequence identification. Microb Genom 2021;7.: 10.1099/mgen.0.000685.

55 Hyatt D, Chen G-L, Locascio PF, Land ML, Larimer FW, Hauser LJ. Prodigal: prokaryotic gene recognition and translation initiation site identification. BMC Bioinformatics 2010;11:119. 10.1186/1471-2105-11-119.

56 Page AJ, Cummins CA, Hunt M, Wong VK, Reuter S, Holden MTG, et al. Roary: rapid large-scale prokaryote pan genome analysis. Bioinformatics 2015;31:3691–3. 10.1093/bioinformatics/btv421.

57 Baba T, Bae T, Schneewind O, Takeuchi F, Hiramatsu K. Genome sequence of Staphylococcus aureus strain Newman and comparative analysis of staphylococcal genomes: polymorphism and evolution of two major pathogenicity islands. J Bacteriol 2008;190:300–10. 10.1128/JB.01000-07.

58 Kläui AJ, Boss R, Graber HU. Characterization and Comparative Analysis of the Staphylococcus aureus Genomic Island vSaβ - an in silico Approach. J Bacteriol 2019. 10.1128/JB.00777-18.

59 Horesh G, Taylor-Brown A, McGimpsey S, Lassalle F, Corander J, Heinz E, et al. Different evolutionary trends form the twilight zone of the bacterial pan-genome. Microb Genom 2021;7:2021.02.15.431222. 10.1099/mgen.0.000670.

60 Read TD, Brunham RC, Shen C, Gill SR, Heidelberg JF, White O, et al. Genome sequences of Chlamydia trachomatis MoPn and Chlamydia pneumoniae AR39. Nucleic Acids Res 2000;28:1397–406.

61 Dobrindt U, Hacker J. Whole genome plasticity in pathogenic bacteria. Curr Opin Microbiol 2001;4:550–7. 10.1016/s1369-5274(00)00250-2.

62 David MZ, Daum RS. Community-associated methicillin-resistant Staphylococcus aureus: epidemiology and clinical consequences of an emerging epidemic. Clin Microbiol Rev 2010;23:616–87. 10.1128/CMR.00081-09.

63 Diekema DJ, Richter SS, Heilmann KP, Dohrn CL, Riahi F, Tendolkar S, et al. Continued emergence of USA300 methicillin-resistant Staphylococcus aureus in the United States: results from a nationwide surveillance study. Infect Control Hosp Epidemiol 2014;35:285–92. 10.1086/675283.

64 Vos M, Eyre-Walker A. Are pangenomes adaptive or not? Nat Microbiol 2017:1576. 10.1038/s41564-017-0067-5.

65 McInerney JO, McNally A, O’Connell MJ. Why prokaryotes have pangenomes. Nature Microbiology 2017;2:17040. 10.1038/nmicrobiol.2017.40.

66 Richardson EJ, Bacigalupe R, Harrison EM, Weinert LA, Lycett S, Vrieling M, et al. Gene exchange drives the ecological success of a multi-host bacterial pathogen. Nat Ecol Evol 2018. 10.1038/s41559-018-0617-0.

67 Mira A, Ochman H, Moran NA. Deletional bias and the evolution of bacterial genomes. Trends Genet 2001;17:589–96.

68 Petit RA, Read TD. Bactopia: a Flexible Pipeline for Complete Analysis of Bacterial Genomes. mSystems 2020;5.: 10.1128/mSystems.00190-20.

69 Di Tommaso P, Chatzou M, Floden EW, Barja PP, Palumbo E, Notredame C. Nextflow enables reproducible computational workflows. Nat Biotechnol 2017;35:316–9. 10.1038/nbt.3820.

70 Seemann T. *snippy: Rapid haploid variant calling and core genome alignment*. 2023. URL: https://github.com/tseemann/snippy (Accessed 25 September 2023).

71 Souvorov A, Agarwala R, Lipman DJ. SKESA: strategic k-mer extension for scrupulous assemblies. Genome Biol 2018;19:153. 10.1186/s13059-018-1540-z.

72 Gurevich A, Saveliev V, Vyahhi N, Tesler G. QUAST: quality assessment tool for genome assemblies. Bioinformatics 2013;29:1072–5. 10.1093/bioinformatics/btt086.

73 Parks DH, Imelfort M, Skennerton CT, Hugenholtz P, Tyson GW. CheckM: assessing the quality of microbial genomes recovered from isolates, single cells, and metagenomes. Genome Res 2015. 10.1101/gr.186072.114.

74 Ondov BD, Treangen TJ, Melsted P, Mallonee AB, Bergman NH, Koren S, et al. Mash: fast genome and metagenome distance estimation using MinHash. Genome Biol 2016;17:132. 10.1186/s13059-016-0997-x.

75 Li H. A statistical framework for SNP calling, mutation discovery, association mapping and population genetical parameter estimation from sequencing data. Bioinformatics 2011;27:2987–93. 10.1093/bioinformatics/btr509.

76 Jain C, Rodriguez-R LM, Phillippy AM, Konstantinidis KT, Aluru S. High throughput ANI analysis of 90K prokaryotic genomes reveals clear species boundaries. Nat Commun 2018;9:5114. 10.1038/s41467-018-07641-9.

77 Buchfink B, Xie C, Huson DH. Fast and sensitive protein alignment using DIAMOND. Nat Methods 2015;12:59–60. 10.1038/nmeth.3176.

78 Page AJ, Taylor B, Delaney AJ, Soares J, Seemann T, Keane JA, et al. SNP-sites: rapid efficient extraction of SNPs from multi-FASTA alignments. Microb Genom 2016;2:e000056. 10.1099/mgen.0.000056.

79 Price MN, Dehal PS, Arkin AP. FastTree 2--approximately maximum-likelihood trees for large alignments. PLoS One 2010;5:e9490.

80 Yu G, Smith DK, Zhu H, Guan Y, Lam TT-Y. Ggtree: An r package for visualization and annotation of phylogenetic trees with their covariates and other associated data. Methods Ecol Evol 2017;8:28–36. 10.1111/2041-210x.12628.

81 Nguyen L-T, Schmidt HA, von Haeseler A, Minh BQ. IQ-TREE: a fast and effective stochastic algorithm for estimating maximum-likelihood phylogenies. Mol Biol Evol 2015;32:268–74. 10.1093/molbev/msu300.

82 Altschul SF, Gish W, Miller W, Myers EW, Lipman DJ. Basic local alignment search tool. J Mol Biol 1990;215:403–10. 10.1016/S0022-2836(05)80360-2.

83 Kassambara A. *rstatix: Pipe-friendly Framework for Basic Statistical Tests in R*. 2023. URL: https://github.com/kassambara/rstatix (Accessed 19 December 2023).

84 Wickham H. Ggplot2. New York, NY: Springer; 2011.

85 drawio: draw.io is a JavaScript, client-side editor for general diagramming and whiteboarding. 2023. URL: https://github.com/jgraph/drawio (Accessed 25 September 2023).

86 Bogart S. *sankeymatic: Make Beautiful Flow Diagrams*. 2023. URL: https://github.com/nowthis/sankeymatic (Accessed 25 September 2023).

87 Jalil M, Quddos F, Anwer F, Nasir S, Rahman A, Alharbi M, et al. Comparative Pan-Genomic Analysis Revealed an Improved Multi-Locus Sequence Typing Scheme for Staphylococcus aureus. Genes 2022;13.: 10.3390/genes13112160.

88 Liu N, Liu D, Li K, Hu S, He Z. Pan-Genome Analysis of Staphylococcus aureus Reveals Key Factors Influencing Genomic Plasticity. Microbiol Spectr 2022;10:e0311722. 10.1128/spectrum.03117-22.

89 Bosi E, Monk JM, Aziz RK, Fondi M, Nizet V, Palsson BØ. Comparative genome-scale modelling of Staphylococcus aureus strains identifies strain-specific metabolic capabilities linked to pathogenicity. Proc Natl Acad Sci U S A 2016. 10.1073/pnas.1523199113.

90 Park S, Jung D, O’Brien B, Ruffini J, Dussault F, Dube-Duquette A, et al. Comparative genomic analysis of Staphylococcus aureus isolates associated with either bovine intramammary infections or human infections demonstrates the importance of restriction-modification systems in host adaptation. Microb Genom 2022;8.: 10.1099/mgen.0.000779.

91 Sassi M, Bronsard J, Pascreau G, Emily M, Donnio P-Y, Revest M, et al. Forecasting Staphylococcus aureus Infections Using Genome-Wide Association Studies, Machine Learning, and Transcriptomic Approaches. mSystems 2022:e0037822. 10.1128/msystems.00378-22.

92 Aanensen DM, Feil EJ, Holden MTG, Dordel J, Yeats CA, Fedosejev A, et al. Whole-Genome Sequencing for Routine Pathogen Surveillance in Public Health: a Population Snapshot of Invasive Staphylococcus aureus in Europe. MBio 2016;7.: 10.1128/mBio.00444-16.

